# Pronounced Genetic Separation Among Varieties of the *Primula cusickiana* Species Complex, a Great Basin Endemic

**DOI:** 10.1101/2021.03.19.436082

**Authors:** Austin Koontz, William D. Pearse, Paul Wolf

## Abstract

Distinguishing between unique species and populations with strong genetic structure is a common challenge in population genetics, especially in fragmented habitats where allopatric speciation may be widespread and distinct groups may be morphologically similar. Such is often the case with species complexes across sky island environments. In these scenarios, biogeography may help to explain the relations between species complex members, and RADseq methods are commonly used to compare closely related species across thousands of genetic loci. Here we use RADseq to clarify the relations between geographically distinct but morphologically similar varieties of the *Primula cusickiana* species complex, and to contextualize past findings of strong genetic structure among populations within varieties. Our genomic analyses demonstrate pronounced separation between isolated populations of this Great Basin endemic, indicating that the current varietal classification of complex members is inaccurate and emphasizing their conservation importance. We discuss how these results correspond to recent biogeographical models used to describe the distribution of other sky island taxa in western North America. Our findings also fit into a wider trend observed for alpine *Primula* species complexes, and we consider how heterostylous breeding systems may be contributing to frequent diversification via allopatric speciation in this genus.

---

A canonical driver of biological diversification is allopatry, whereby geographic barriers lead to population isolation and, eventually, speciation. Sky islands are places where sharp changes in elevation lead to pronounced ecological differences over relatively short distances, providing the types of barriers required for allopatric speciation to take place. Historically, climatic fluctuations have determined the presence and distribution of sky island environments for mountain ranges across the world, and this in turn is reflected by the genetic patterns seen in montane species today (Hewitt 2000). However, in this biogeographic context, distinguishing between closely related species and genetically structured populations may prove challenging (Huang 2020), especially if similar niches across mountain ranges maintain phenotypic similarities (e.g. Yang et al. 2019). Additionally, in the short-term, genetic patterns will be influenced by particular aspects of a species’ biology, such as dispersal and breeding systems, which may facilitate or hinder reproductive isolation between genetically distinct entities. Here, we examine the genomic relations between the sky island populations of members of the *Primula cusickiana* species complex, a group of plants endemic to the Great Basin region of the western United States.

The *P. cusickiana* species complex is a group of herbaceous, perennial plants that fall within the Parryi section of *Primula*. The morphological differences between the four complex varieties—*maguirei, cusickiana, nevadensis*, and *domensis* (see Fig. 1)*—*are subtle: *maguirei* (Williams 1936) and *cusickiana* (Gray 1888) are entirely glabrous, and distinguished from one another by relative calyx length, while in *nevadensis* (Holmgren 1967) and *domensis* (Kass and Welsh 1985), plants are pubescent and have slightly different corolla tube lengths (Holmgren and Kelso 2001; Holmgren et al. 2005). Despite these subtle differences, varieties *cusickiana, nevadensis*, and *maguirei* were originally classified as separate species, based on ecological traits and distinct geographic ranges. The discovery and publication of *P. domensis* in 1985, along with the continued collection of the other varieties, began to cast doubt on the species distinction for each complex member. A 2001 review determined that the morphological differences were insufficient for species classification, and subsumed each species to the level of variety (Holmgren and Kelso 2001).

**Fig 1.**
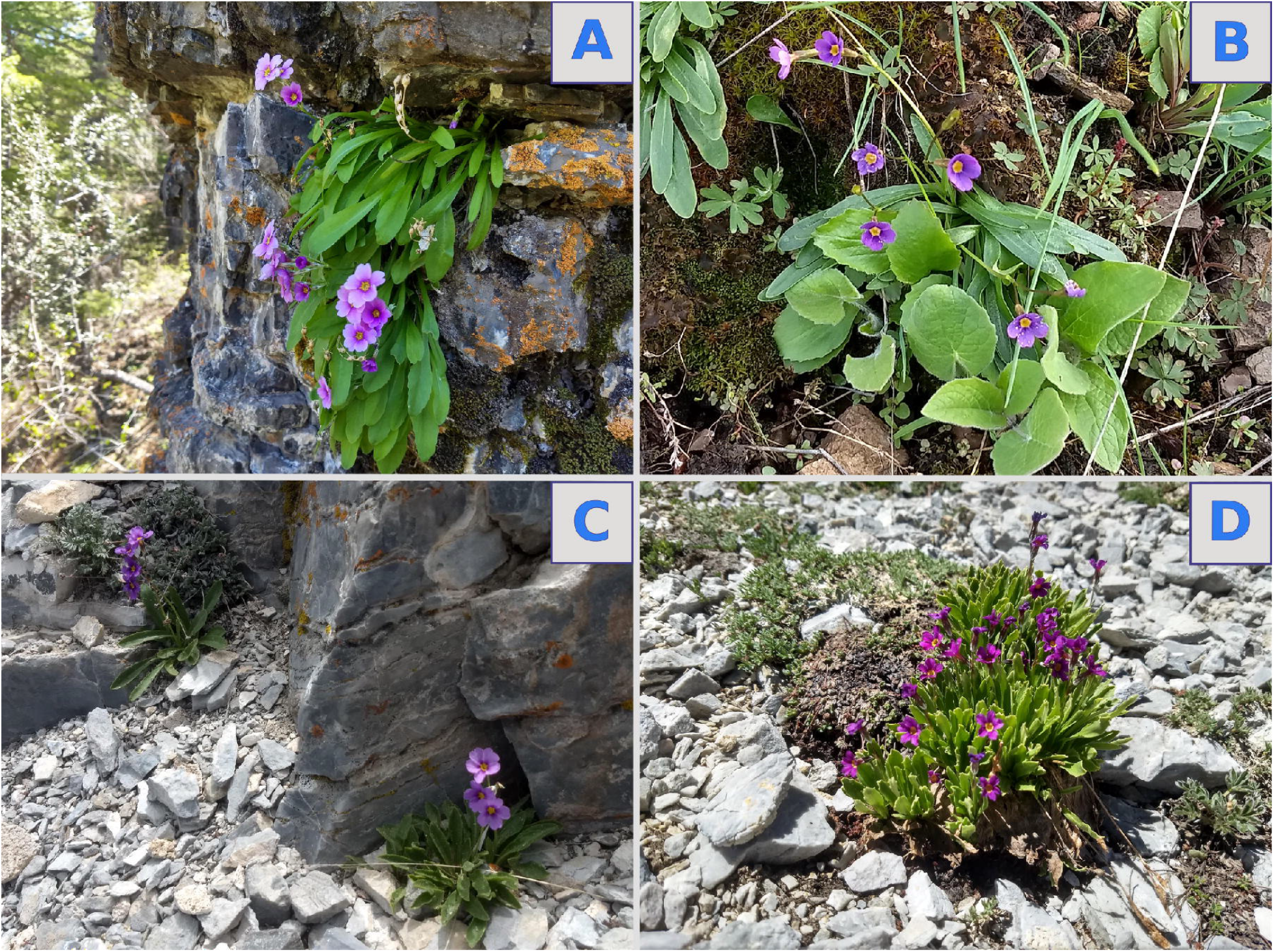
Four members of the *Primula cusickiana* species complex: (A) *maguirei*, in Right Hand Fork of Logan Canyon; (B) *cusickiana*, near Cougar Point in Jarbidge, Nevada; (C) *domensis*, at Notch Peak in the House Range, Utah; (D) *nevadensis*, on Mount Washington in the Snake Range (Great Basin National Park), in Nevada.

At the time of this shift, no genetic data was available to justify classification at the variety level. However, a 1997 analysis of variety *maguirei* used allozyme marker genes to uncover a significant degree of genetic structure between the relatively proximate (∼10 km) populations (Wolf and Sinclair 1997) within this one taxon. A later analysis of the same populations using amplified fragment length polymorphism (AFLP) loci confirmed this finding, and found similar levels of polymorphism between the upper and lower canyon groups, suggesting this genetic structure is not the result of a past bottleneck event (Bjerregaard and Wolf 2004). A further analysis of AFLP and chloroplast DNA from the *Primula* section Parryi showed *maguirei* and the other *P. cusickiana* complex members as being monophyletic, but relationships within the complex were incongruent, with only weak support of a clade containing *nevadensis* and *domensis* being sister to a clade made up of *maguirei* and *cusickiana* (Kelso et al. 2009). To better resolve the relationships between varieties, the authors suggested an analysis utilizing more populations from across the range of this species complex. Restriction-site associated sequencing (RADseq) technologies available today, with their ability to generate reads over many sequence regions of closely related individuals, are well-suited to provide the data required for such an analysis.

In addition to clarifying the genetic relations between geographically distinct varieties, a more detailed analysis of the *P. cusickiana* species complex can meaningfully contribute to ongoing conservation efforts. Variety *maguirei* was listed as Threatened in 1985, due to its unique habitat in Logan Canyon and threats of habitat loss due to development (Fish and Wildlife Service 1985). Given the strong genetic structure between *maguirei*’s populations, either population may be more closely related to populations of a different complex variety than the neighboring Logan Canyon population—a finding which would have significant implications for the protection of this variety. More broadly, an understanding of the genomic relations at the species complex level will determine whether the varietal classification properly reflects the extent of genomic divergence of each complex member, and thus the extent of unique evolutionary history. This understanding can direct management of the narrow-range endemics included in this species complex—such as *maguirei*, but also *nevadensis* and *domensis*—and also inform the identification of potential evolutionary significant units (Coates et al. 2018).

We sought to clarify the relatedness of *P. cusickiana* complex members by using a RADseq approach to genotype all four varieties located at distinct populations scattered throughout the Great Basin. In addition to contextualizing the genetic structure between the upper and lower Logan Canyon *maguirei* populations, this analysis provides insights into the biogeographic history of this species complex, and could have important conservation implications for this rare endemic plant.

## Materials and Methods

### Sampling

All *P. cusickiana* species complex samples were gathered in the field, along with samples of *P. parryi* (Gray 1888), which was used as an outgroup in genetic analyses. Populations and their respective flowering times were determined using herbarium specimens, and collection sites were selected to maximize the geographic distribution of each variety. At each population location, an individual plant was removed as completely as possible as a voucher specimen. For DNA samples, two leaves from each of ten plants were removed and placed in labeled paper envelopes, which were stored on silica crystals to keep samples dry. Vouchers were deposited at the Intermountain Herbarium (UTC); *P. cusickiana* var. *nevadensis* voucher specimens collected from Mt. Washington were additionally deposited at the Great Basin National Park herbarium.

Because past research has shown variable relations between *P. capillaris* (Holmgren and Holmgren 1974) and the *P. cusickiana* species complex (Kelso et al. 2009), we also tried to collect *P. capillaris* in the field. However, we were unable to locate any *P. capillaris* individuals in the Ruby Mountains: at one location suggested by past herbaria data, a population of *P. parryi* was found instead. To compensate, two *P. capillaris* samples were sourced from herbaria (see Appendix I).

Leaf tissue from 89 samples—87 silica-dried field collections representing all samples sites, and two herbarium specimens of *P. capillaris*—were placed into QIGAEN Collection Microtubes (catalog number 19560) and sent to University of Wisconsin-Madison Biotechnology Center, for DNA extraction, library prep, and DNA sequencing (described below). Seven replicate samples were also included to assess the quality of sequencing results, and were distributed across all four *P. cusickiana* varieties, as well as *P. parryi*.

### DNA Extraction

DNA was extracted using the QIAGEN Dneasy mericon 96 QIAcube HT Kit. DNA was quantified using the Quant-iT^™^ PicoGreenR^©^ dsDNA kit (Life Technologies, Grand Island, New York).

### Library Prep and Sequencing

Libraries were prepared following Elshire et al. 2011. *ApekI* (New England Biolabs, Ipswich, Massachusetts) was used to digest 100 ng of DNA. Following digestion, Illumina adapter barcodes were ligated onto DNA fragments using T4 ligase (New England Biolabs, Ipswich, Massachusetts). Size selection was run on a PippinHT (Sage Science, Inc., Beverly, Massachusetts) to subset samples down to 300—450 bp fragments, after which samples were purified using a SPRI bead cleanup. To generate quantities required for sequencing, adapter-ligated samples were pooled and then amplified, and a post-amplification SPRI bead cleanup step was run to remove adapter dimers. Final library qualities were assessed using the Agilent 2100 Bioanalyzer and High Sensitivity Chip (Agilent Technologies, Inc., Santa Clara, California), and concentrations were determined using the Qubit^©^ dsDNA HS Assay Kit (Life Technologies, Grand Island, New York). Libraries were sequenced on an Illumina NovaSeq 6000 2×150.

### Data Processing

Raw FASTQ data files were demultiplexed and processed using steps 1—7 of the *ipyrad* software, version 0.9.31 (Eaton and Overcast 2020). Single nucleotide polymorphisms (SNPs) recognized by *ipyrad* were used as the basis for variation between individuals for downstream analyses, and libraries were assembled *de novo*. All *ipyrad* and STRUCTURE parameter files, as well as R scripts used for analysis and data visualization, can be found on GitHub (github.com/akoontz11/Primula/) and in the Supplementary Materials (SupplementalMaterials1.zip). Raw, demultiplexed sequencing data can be accessed on the NCBI Sequence Read Archive (SRA; accession number PRJNA705310).

#### Complex-Wide Genomic Survey

For our complex-wide genomic survey, we ran *ipyrad* twice: we used the results from our initial run to confirm sequencing consistency for replicate samples, and to identify samples with low coverage. For both runs, demultiplexed sequences were paired and merged, and low quality bases, adapters, and primers were filtered prior to SNP calling. Default values were used for the *ipyrad* parameters in these steps, as well as for the clustering threshold (clust_threshold; 0.85) and minimum sequencing depth (mindepth_statistical; 6) parameters.

For our initial run, we specified a minimum number of samples per locus (min_samples_locus) parameter of 10, in order to obtain loci shared between two to three sample locations for any variety. Using the results from this run, we used the Python script vcf2Jaccard.py to compare samples with replicates by calculating the mean Jaccard similarity coefficients between all samples. We found that all replicates matched highly with their corresponding samples (Fig. S1).

After merging replicates and removing low coverage (generally, less than 30 loci in the final assembly) samples from the dataset, 82 of our 87 original samples remained for our complex-wide analysis. We reran *ipyrad* (steps 1-7) using these 82 samples to select for loci specific to this subset. We used a min_samples_locus parameter of 32 for this second run, to match the ratio of minimum samples per locus used in our initial run; ipyrad default values were used otherwise. Because very low numbers of loci were retrieved for both herbarium specimens of P. capillaris (possibly due to the age of these specimens), we were unable to include *capillaris* in downstream clustering analyses.

#### Variety Specific Clustering

In addition to our complex-wide survey, we were interested in exploring population structure within variety maguirei which could not be resolved using genetic loci shared across all species complex members. To do so, we ran ipyrad on just the 18 maguirei samples used in our complex-wide survey. Because five samples from each of the upper Logan canyon sampling sites were included in our ipyrad assembly, we specified a min_samples_locus parameter of 5; ipyrad default parameter values were used otherwise.

### Population Analyses

#### Structure

To visualize relations between complex members across their geographic range, and to determine the number of identifiable genetic clusters within the complex, we used the program STRUCTURE version 2.3 (Pritchard et al. 2000). STRUCTURE uses Bayesian clustering analysis to probabilistically assign individuals to one or more of K source populations, where the loci within each population are assumed to be in Hardy-Weinberg proportions and linkage equilibrium. For all STRUCTURE runs, we used a burnin length of 50,000, and 100,000 MCMC reps after burnin. For our complex-wide survey, we ran STRUCTURE for K values of 2—16, with 50 replicates per K value. For our *maguirei*-only analyses, we ran STRUCTURE for K values of 2—6, with 50 replicates per K value. We used the CLUMPAK server (Kopelman et al. 2015) to summarize results across replicates for each K value, and to build STRUCTURE plots.

For all of our STRUCTURE analyses, we ran the Evanno et al. (2005) method (which identifies the greatest ΔK value) and the method described in the STRUCTURE manual (Pritchard et al. 2000, which identifies the K value with the greatest likelihood) to determine an “optimal” K value. Given the difficulties in inferring an unambiguous number of genetic clusters from any given set of populations (Novembre 2016; Pritchard et al. 2000), we also examined STRUCTURE outputs within a range of K values, to determine which value of source populations best illustrated divisions within the species complex.

#### Discriminant Analysis of Principal Components

In addition to STRUCTURE, we analyzed the results of our complex-wide survey using Discriminant Analysis of Principal Components (DAPC; Jombart et al. 2010) in the package *adegenet* in R version 3.6.3 (R Core Team, 2020). DAPC is a statistical technique designed to accommodate the size of genomic data sets and capable of differentiating within-group variation from between-group variation. SNP data is first transformed using a principal components analysis (PCA), and then k-means clustering is run to generate models and likelihoods corresponding to each number of population clusters. The best-fitting model, and so the best-supported number of populations, is assessed using the models’ Bayesian Information Criterion (BIC) scores. We chose to utilize DAPC in addition to STRUCTURE to visualize population clusters in a PCA format, and to determine whether the supported number of populations was congruent between methods, indicating a more robust determination of the number of species contained within the complex (Carstens et al. 2013).

#### F_ST_ Estimates

Because we wanted to measure the extent of genetic variance within the groups analyzed, we used the VCFtools software (Danecek et al. 2011) to generate weighted FST estimates (Weir and Cockerham 1984). We generated an FST estimate for our complex-wide analysis (across all populations of all *P. cusickiana* varieties) as well as for the samples included in our variety *maguirei*-only analysis.

## Results

### Complex-Wide Genomic Survey

We retrieved, on average, 2.04 x 106 reads per sample, and our complex-wide *ipyrad* run identified 1,277 loci that were used in our subsequent STRUCTURE analysis. Using the Evanno et al. (2005) method yielded an optimal K value of K = 5; using the method described in the STRUCTURE manual (Pritchard et al. 2000) identified the K value with the greatest likelihood as K = 14. Based on our visualization of the STRUCTURE results for values ranging from K = 2—16 (Figs. S2 - S4), we determined K = 7 to be the most biologically relevant K value (Fig. 2). At this level of source populations, varieties *domensis* and *maguirei* are clearly delineated, variety *nevadensis* shows distinctions between its two populations, and variety *cusickiana* is split into three groups composed of populations from the Snake River Plain in Idaho (SRP), Nevada (Jarbidge), and Oregon (Owyhee). Since higher K values emphasize the divisions seen at this level, and further subdivide isolated populations of varieties *cusickiana* and *nevadensis*, K = 7 is a conservative estimate which reflects the strong divisions within this complex while allowing for further distinctions between unique populations to be made in light of more evidence in the future.

**Fig 2.**
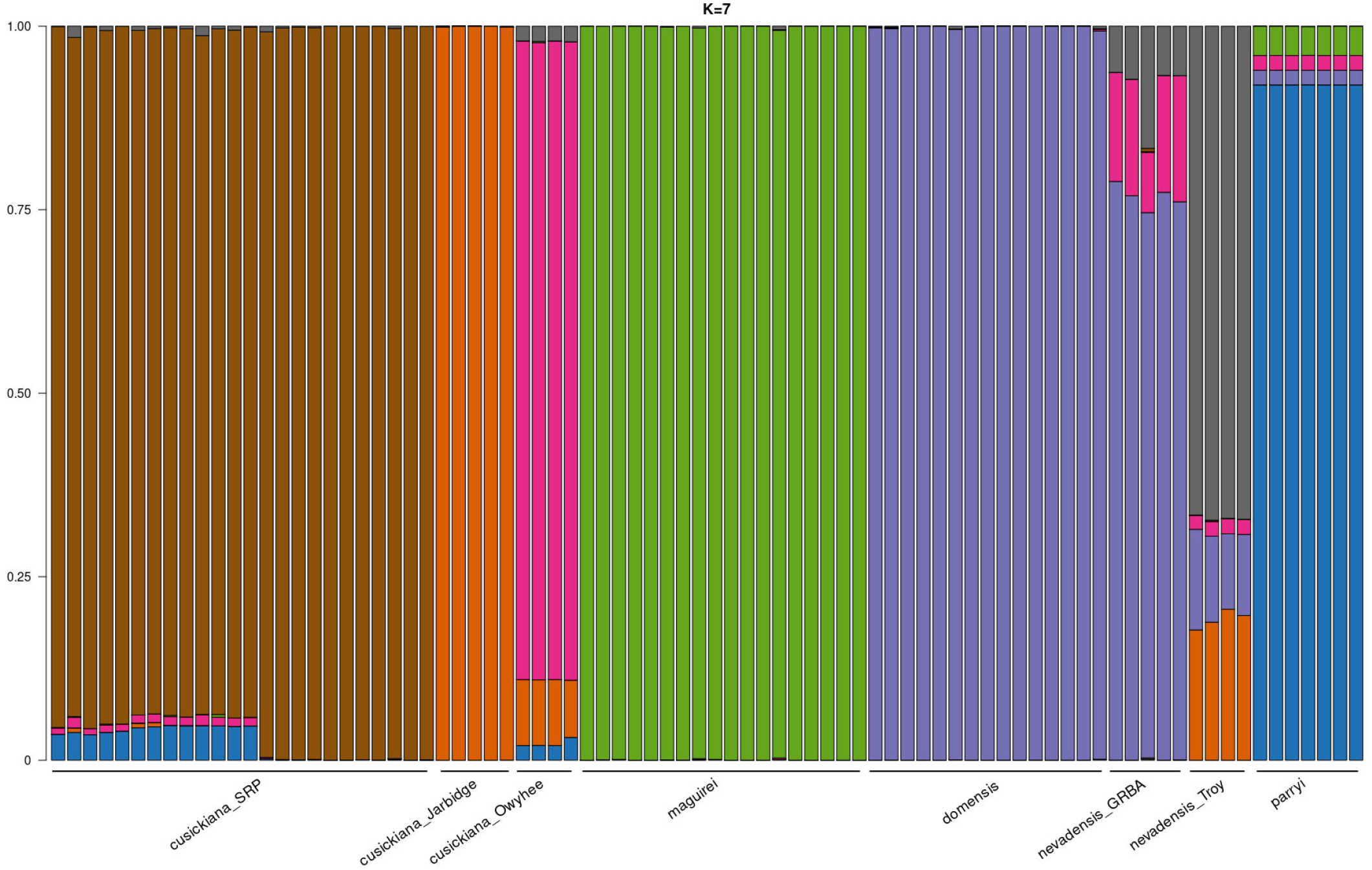
Sample STRUCTURE plots at K = 7. At this level of clustering, divisions between isolated populations of variety *cusickiana* in Idaho (Snake River Plain, or SRP), Nevada (Jarbidge), and Oregon (Owyhee) are clearly shown. Similarities between populations of variety *nevadensis* in Great Basin National Park (GRBA) and domensis are shown, while populations of *nevadensis* further south in the Grant Range (Troy) are more distinct.

Our DAPC analysis revealed that the greatest supported number of clusters (i.e. the value with the lowest BIC score) was eleven (data not shown)—a value incongruent with our STRUCTURE results, suggesting that boundaries within this complex are elaborate. However, at this level of genetic clusters, several groups were quite small (consisting of only one or two samples), and groupings were incoherent within the spatial distribution of populations. To provide a clearer comparison to our STRUCTURE results, and to examine relations strictly within the species complex, we removed *P. parryi* outgroup samples from our dataset (because these were separate from all species complex samples in preliminary analyses) and ran our DAPC with a specification of six clusters (Fig. 3). At this level of clustering, the population of *nevadensis* in the Snake Range of Great Basin National Park (GRBA) is shown as a unique cluster, while the *nevadensis* population further south in the Grant Range groups with the *cusickiana* population sampled from Oregon (Owyhee). Variety *domensis* is a unique cluster which groups closely to both of these. Thus, while neither our STRUCTURE analysis nor our DAPC point to an unambiguous number of “true” genetic clusters, both suggest that the current varietal classification is inexact. The extreme level of divergence between the sky island populations in this species complex is reflected not only in our clustering analyses, but also in our relatively large FST estimate across all complex populations, which was 0.72. Figure 4 illustrates proportions of sample membership to clusters based on our STRUCTURE analysis at K=7 for all populations in their geographic context across the Great Basin.

**Fig 3.**
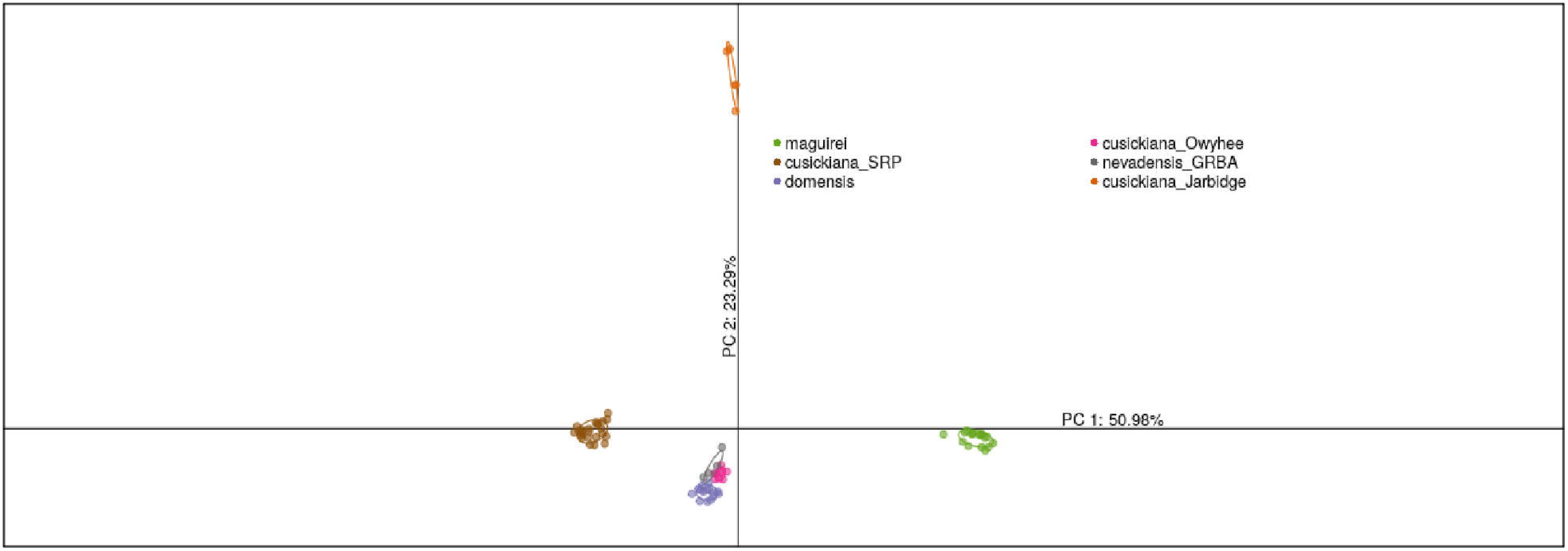
DAPC of only *P. cusickiana* complex samples with number of genetic clusters specified at 6; percentage of total variance for each PC axis shown. Similar to STRUCTURE results at K = 7, this analysis shows all *maguirei* populations as distinct from all other complex populations. Populations of varieties *domensis* and *nevadensis* group closely with *cusickiana* population from Oregon (“cusickiana_Owyhee”).

**Fig 4.**
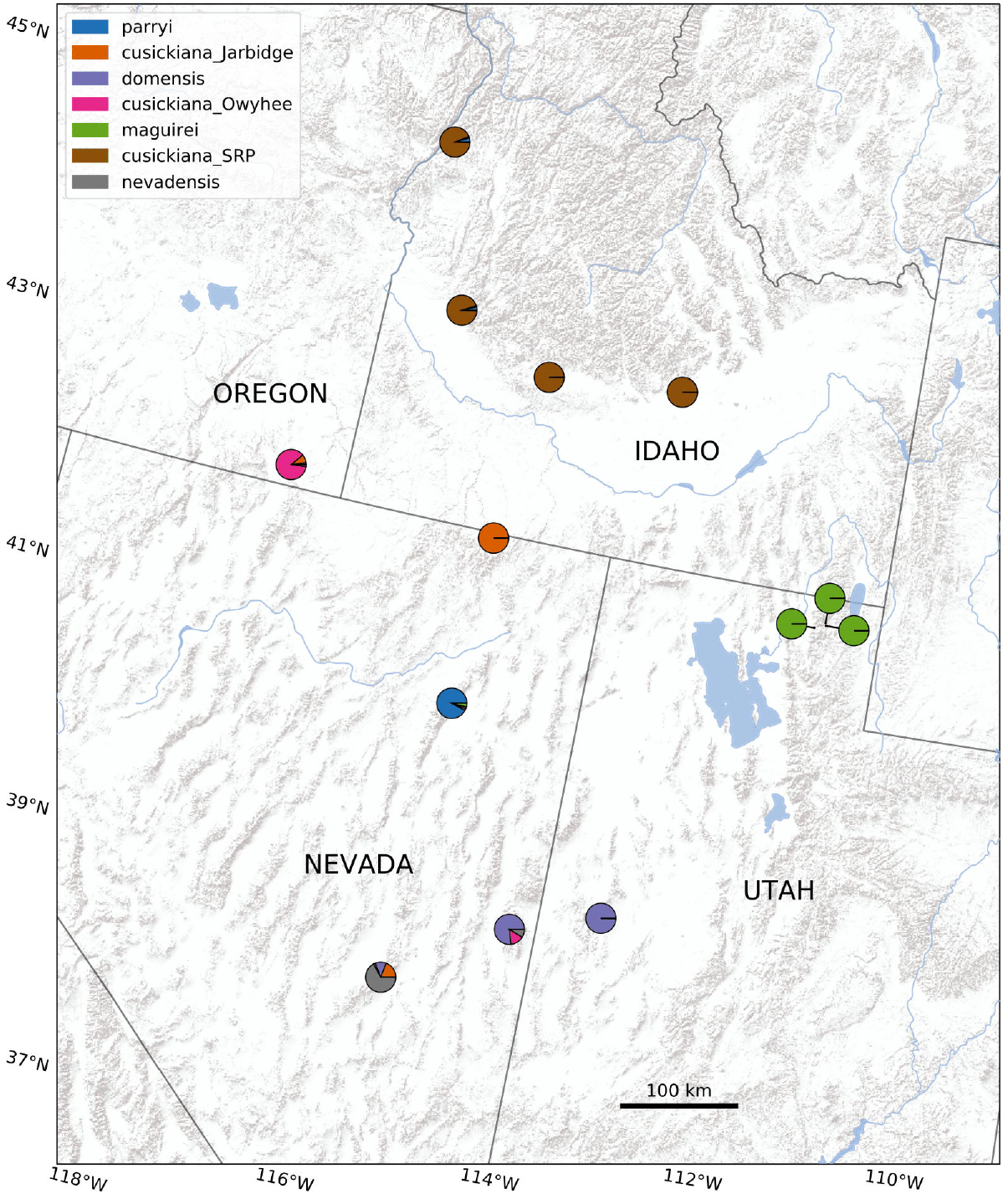
Map of sample locations with cluster membership. Sampling locations are represented by pie charts indicating percentage of population membership to clusters determined at K = 7 STRUCTURE clustering threshold. With exception to *nevadensis*, most samples fall almost entirely within a specified cluster.

### Variety Specific Clustering

In our complex-wide analysis, all *maguirei* samples grouped as a single cluster, distinct from all other populations of all other varieties, indicating that neither Logan Canyon population is more closely related to any populations of another variety. Even at values of K = 16, the upper and lower Logan Canyon populations of *maguirei* were not resolved from one another.

However, reducing our sample set to only *maguirei* samples allowed us to retain loci informative to this variety but unshared with other complex member populations. Our *maguirei*-only *ipyrad* run generated an assembly with 68,492 loci, indicating a large number of loci specific to *maguirei* and not shared with the wider species complex. To speed up processing times, we ran STRUCTURE on a 17,988 loci subset of *maguirei*-specific markers. Using the CLUMPAK server, we found optimal K values of K = 4 (using the Evanno method) and K = 3 (using the likelihood method described in the STRUCTURE manual). Figure 5 shows the STRUCTURE plot at K = 3, which resolves similar groupings of *maguirei* populations supported in Bjerregaard and Wolf (2004), and the distinctions between upper and lower canyon populations. We also estimated an F_ST_ value of 0.33 among these three populations, which is comparable to previous estimates in Bjerregaard and Wolf (2004).

**Fig 5.**
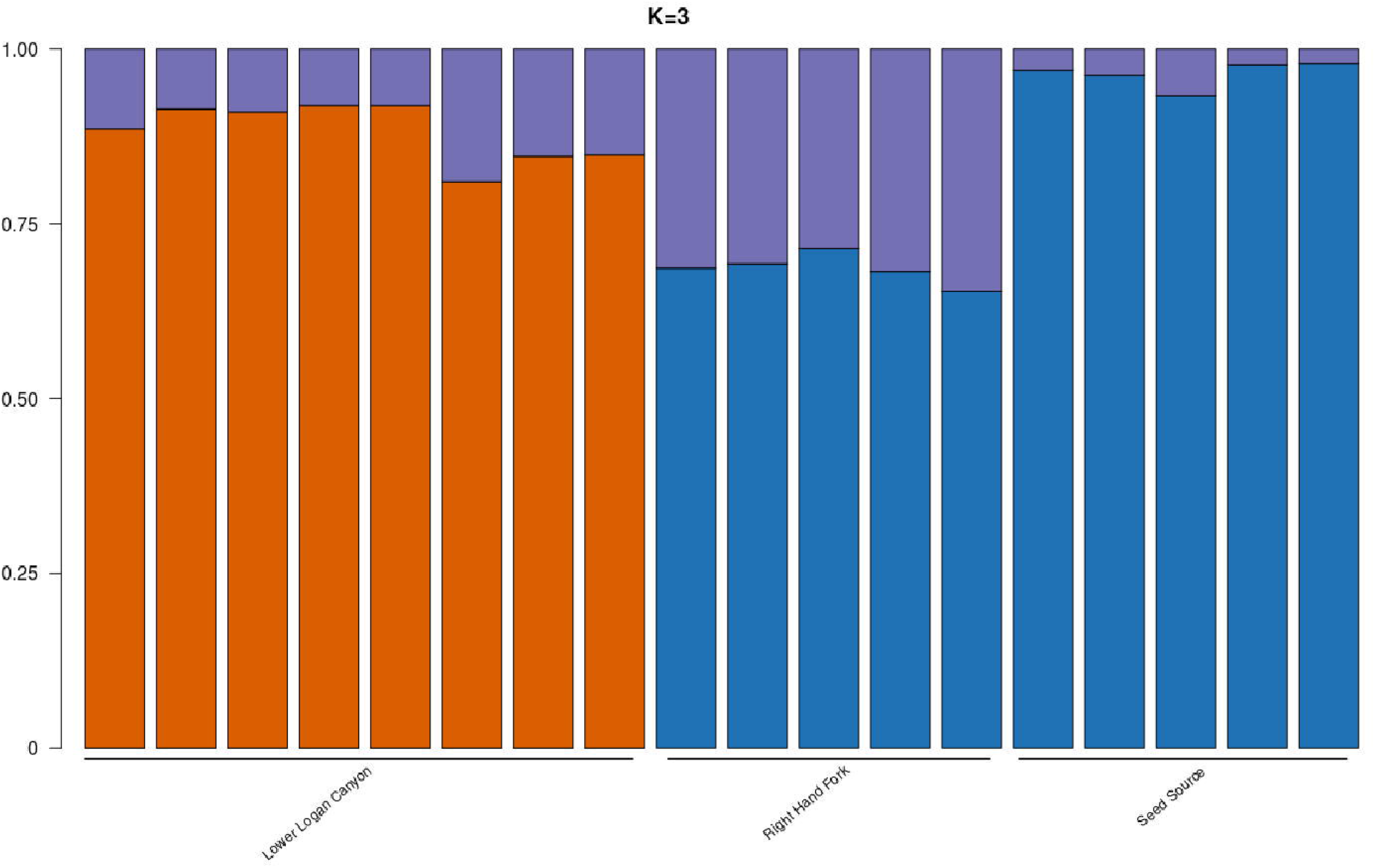
STRUCTURE plot for *maguirei* samples at a clustering threshold of K = 3. While *maguirei* clustered together in the complex-wide analysis, our *maguirei*-only analysis was able to reveal the Logan Canyon population divisions illustrated in past studies.

## Discussion

Analysis of RADseq data from *Primula cusickiana* complex members demonstrates that the disjunct geographical distribution of populations across the Great Basin is reflected by pronounced genomic divergences. While the results of our clustering analyses coincide with the current varietal classifications, there are notable exceptions. Distinctions between isolated populations within varieties, as well as similarities between neighboring populations of different varieties, can be observed in our STRUCTURE plots for low K values (i.e. ranging from 2—6; see FIGS. S2-S4). For instance, we found Mt. Washington *nevadensis* populations to be admixed, with segments coming from *domensis* to the east and (to a lesser extent) Grant Range *nevadensis* populations to the south. This is in accordance with analysis of AFLP and chloroplast DNA from the *Primula* section Parryi, which found these two varieties to be extremely close (Kelso et al. 2009).

Our results also suggest a more nuanced understanding of variety *cusickiana*. Populations of this variety are split into distinct genomic clusters in our analysis, with Jarbidge (Nevada) and Owyhee (Oregon) populations appearing unique from each other and the remaining Snake River Plain (SRP) populations in Idaho. That these distinctions are seen in both our STRUCTURE and DAPC analyses imply the robustness of this result. Given the relatively wide distribution of this variety (growing in moist soils at lower elevations than other complex members), our findings of genomic divergence between its populations is noteworthy, and support past evidence of phenotypic differences in different portions of its range. For instance, past morphological research of Idaho *cusickiana* populations has suggested dividing this taxa into three unique species (Mansfield 1993), with Owyhee populations being classified as *P. wilcoxiana*.

The separation between populations within variety *cusickiana*, as well as our support of past findings of significant genetic distances between the proximate populations of variety *maguirei*, underscore our discovery of profound genomic divergences between all members of this species complex, despite their distribution over a relatively small geographic area. This trend is reflected not only in our clustering analyses, but also in our weighted F_ST_ estimate of 0.72 across complex populations—a high value compared to similar estimates for other plant taxa (for instance, the mean F_ST_for plant taxa in a meta-analysis by Leinonen et al. 2008 was calculated to be 0.24). Our results therefore support the historical designation of species for these complex members, rather than variety. Below, we consider how two phenomena—biogeographical trends in the Great Basin, and reproductive traits specific to *Primula*—may contribute to the significant divergence of these populations into distinct genomic groups.

### Great Basin Sky Island Biogeography

Members of the *P. cusickiana* complex are found at relatively high elevations throughout the Great Basin. Many of these are sky island locations associated with strong ecological shifts as habitat transitions from lower sagebrush steppe to cooler, more forested regions dominated by pinyon and juniper. Now separated by arid basins due to climatic warming in the Holocene, these sky islands are understood to be the fragmented remnants of a continuous region of cool, moist habitat which once extended across the Great Basin (Thompson and Mead 1982). This has led to their characterization as refugia for various taxa—particularly mammals (Brown 1971; Badgley et al. 2014), but also butterflies (Boggs and Murphy 1997) and plants (Harper et al. 1978; Nowak et al. 1994; Charlet 2007). Additionally, in conjunction with climatic niche preferences, complex varieties *maguirei, domensis*, and *nevadensis* are found on the cliffs and crevices of exclusively limestone substrates. While it’s unclear whether these habitats are tied to mineral or pH constraints, or simply reflect preferences for moisture-retentive substrates, edaphic heterogeneity is known to contribute to plant speciation and biodiversity, both globally (Hulshof and Spasojevic 2020) and within the Great Basin (e.g. de Queiroz et al. 2012). Therefore, allopatry across relatively similar climatic and edaphic niches seems to contribute to the genomic divergences in *P. cusickiana*’s populations—a trend observed in other sections of Primulaceae, as well (Boucher et al. 2016).

However, it has also been noted that many species distribution patterns among Great Basin mountaintops do not follow a strictly island biogeographical model (Lawlor 1998), in that neither island surface area nor proximity to “mainland” source populations (typically identified as the western Sierra Nevadas or eastern Rocky Mountains) is predictive of species abundance (Fleishman et al. 2001). And in some taxa, there is evidence for regular, modern dispersal between Great Basin ranges (Floyd et al. 2005). An alternative scenario is that this complex has followed what has been described as an “expanding-contracting archipelago” (ECA) model, in response to Quaternary glacial cycles (DeChaine and Martin 2005a). The ECA model has been used to describe the divergence between Rocky Mountain sky island plant taxa (Dechaine and Martin 2005b; Hodel et al. 2021), and provides a framework for explaining the genetic structure observed between isolated montane populations on a broad spatial scale. In this model, populations are assumed to become fragmented as they contract up-slope during warmer interglacials; during glacial periods, populations expand down-slope as moist, cool habitat becomes widespread, leading to hybrid zones and possible admixture. Given the degree of fragmentation between *P. cusickiana*’s populations in today’s climate (which resembles past interglacial periods), and the admixture between the relatively proximate populations of varieties *domensis* and *nevadensis* revealed in our analysis, this model offers a viable explanation for the trends observed in this species complex. In addition to these biogeographic patterns, the evolution of *P. cusickiana*’s disjunct populations is simultaneously influenced on a finer spatial scale by aspects particular to this species’ biology.

### Speciation and Heterostyly in Primula

Recent research has shown several different alpine Primula species complexes to contain previously undescribed cryptic species, in China (Huang et al. 2019; Ren et al. 2020) and in Europe (Schorr et al. 2013; Theodoridis et al. 2019). Our findings on the *P. cusickiana* species complex resonate with these trends, and raise the question of what unique traits *Primula* possesses which might cause such frequent diversification via allopatric speciation. The authors of a study examining the P. *merrilliana* species complex in China (He et al. 2021) argue that heterostyly—a widespread breeding system in angiosperms to promote outcrossing—may be a driving force leading to speciation. In heterostyly, “pin” and “thrum” floral morphologies prevent self-fertilization via insect pollination (Darwin 1897), and are associated with a sporophytic-incompatibility system which follows a Mendelian pattern of inheritance (Li et al. 2016). In *P. merrilliana*, the efficacy and prevalence of heterostyly and self-incompatibility varies across populations, which has possibly led to the divergence between distylous and homostylous populations and, ultimately, speciation.

While the presence of heterostyly has been observed in *nevadensis* (Holmgren 1967) and in populations of *cusickiana* and *domensis* (pers. obs.), the extent of distyly in a population has only been well documented in *maguirei*, who’s upper and lower canyon populations have a pin:thrum morphology ratio of about 1:1 (Davidson et al. 2014). This implies that in scenarios of legitimate xenogamy, in which morphs of one type only mate with morphs of the opposite type, only half of the total population is available as a potential mate for any distylous individual. While this reduction in effective population size would seem to increase the strength of genetic drift, and possibly the fixation of deleterious alleles, these negative effects are potentially counterbalanced by the genetic advantages of outcrossing. This net benefit of heterostyly is supported by findings in de Vos et al. (2014), in which phylogenetic techniques were used to demonstrate that the presence of heterostyly in *Primula* leads to greater diversification via decreased extinction, in the long-term, compared to non-heterostylous clades of Primulaceae. Simultaneously, the loss of heterostyly and subsequent self-compatibility may lead to rapid speciation in the short-term. Observation of distylous morph ratios in other *P. cusickiana* varieties and populations, and changes in these ratios between proximate populations, would help to determine if these dynamics are driving the divergences we see at the species complex level.

## Conclusion

The results of our genomic survey of *Primula cusickiana* fit into a wider trend demonstrating abundant allopatric speciation despite little niche divergence in other alpine *Primula* species complexes. Our findings support the historical classification of each of these complex members as unique species, rather than the varietal classification taken in Holmgren and Kelso (2001). Furthermore, these results warrant a more detailed understanding of the isolated and genetically unique populations in this complex (such *as cusickiana* populations in Nevada and Oregon), and of the admixture observed in the populations of variety *nevadensis*. Similarly, updated morphological comparisons between varieties, as well as observations into the levels of heterostyly in disjunct populations, would offer a clearer understanding of the mechanisms of speciation occurring within this complex. Finally, the endemic species with narrow niches included in this study, such as *P. cusickiana* var. *maguirei*, but also *nevadensis, domensis*, and the sister species P. capillaris, warrant concern of extinction, and more work needs to be done to better understand the breeding limitations faced by each of these taxa and what can be done to ensure their survival in an increasingly arid Great Basin.

## Acknowledgments

his research was supported by funds provided by the Margaret Williams Research Grant offered by the Native Nevada Plant Society, the Lawrence Piette Graduate Scholarship offered by the USU College of Science, the Dr. Ivan J. Palmblad Graduate Research Award offered by the USU Department of Biology, and the Graduate Research Award offered by the USU Ecology Center. WDP and the Pearse Lab are funded by NSF EF-1802605, NSF ABI-1759965, and UKRI-NERC NE/V009710/1.

The authors would like to thank Dr. Carol Rowe for her help with the genetic analyses, her Python script used to calculate Jaccard similarities from SNP data, and her Python script for creating the map. Thanks also to Dr. Barbara Ertter and especially Dr. Don Mansfield at the College of Idaho, for their assistance in collecting variety *cusickiana*. Trish Winn and Jennifer Lewihnsohn in the United States Forest Service provided the collection permit for variety *maguirei* from the Uinta-Wasatch-Cache National Forest; Todd Stefanic and Gretchen Baker with the National Park Service coordinated collection from Craters of the Moon National Monument and Great Basin National Park, respectively. Michael Piep, Elizabeth Makings, and Jerry Tiehm allowed for *Primula* specimens to be sampled from the Intermountain Herbarium. Arizona State Vascular Plant Herbarium, and University of Nevada, Reno Herbarium, respectively. Thanks also to Kris Valles at the Intermountain Herbarium. Dr. Leila Shultz assisted with identification of the Owyhee *cusickiana* population. The authors would also like to thank Jean Howerton, Buddy Smith, and Noel and Pat Holmgren. Finally, all collections were made on the ancestral lands of the Western Shoshone, Eastern Shoshone, Shoshone-Bannock, Southern Paiute, Goshute, and Nez Perce Native American tribes.

## Author Contributions

AK determined sample locations, performed the majority of sample collection, and ran genetic analyses. WDP contributed to study design and assisted with genetic analyses and manuscript writing. PW guided study design and assisted with genetic analyses, sample collection, and manuscript writing.

## Appendix 1

Voucher specimens. Order of data is as follows: Species, Voucher, Herbarium. Institutional barcodes or accession numbers are included as parenthetical values following the voucher, when available.

### Ingroup

*Primula cusickiana* var. *cusickiana*, 25330978, Intermountain Herbarium; *Primula cusickiana* var. *cusickiana*, 25330990, Intermountain Herbarium; *Primula cusickiana* var. *cusickiana*, 25331045, Intermountain Herbarium; *Primula cusickiana* var. *cusickiana*, 25331062, Intermountain Herbarium; *Primula cusickiana* var. *cusickiana*, 25331021, Intermountain Herbarium; *Primula cusickiana* var. *cusickiana*, 25331015, Intermountain Herbarium; *Primula cusickiana* var. *cusickiana*, 25331018, Intermountain Herbarium; *Primula cusickiana* var. *cusickiana*, 25331034, Intermountain Herbarium; *Primula cusickiana* var. *cusickiana*, 25331004, Intermountain Herbarium; *Primula cusickiana* var. *cusickiana*, 25330994, Intermountain Herbarium; *Primula cusickiana* var. *cusickiana*, 25330991, Intermountain Herbarium; *Primula cusickiana* var. *maguirei*, 25331026, Intermountain Herbarium; *Primula cusickiana* var. *maguirei*, 25331039, Intermountain Herbarium; *Primula cusickiana* var. *maguire*i, 25331041, Intermountain Herbarium; *Primula cusickiana* var. *nevadensis*, 25331101, Intermountain Herbarium; *Primula cusickiana* var. *nevadensis*, 25331106, Intermountain Herbarium; *Primula cusickiana* var. *nevadensis*, 25331092, Intermountain Herbarium; *Primula cusickiana* var. *domensis*, 25331066, Intermountain Herbarium; *Primula cusickiana* var. *domensis*, 25331070, Intermountain Herbarium; *Primula cusickiana* var. *domensis*, 25331077, Intermountain Herbarium; *Primula cusickiana* var. *domensis*, 25331083, Intermountain Herbarium;

### Outgroups

*Primula capillaris*, 770850 (ASU0020421), Arizona State University Vascular Plant Herbarium; *Primula capillaris*, 3025822 (UTC00138833), Intermountain Herbarium; *Primula parryi*, 25331110, Intermountain Herbarium; *Primula parryi*, 25331112, Intermountain Herbarium

---

**Fig S1.**
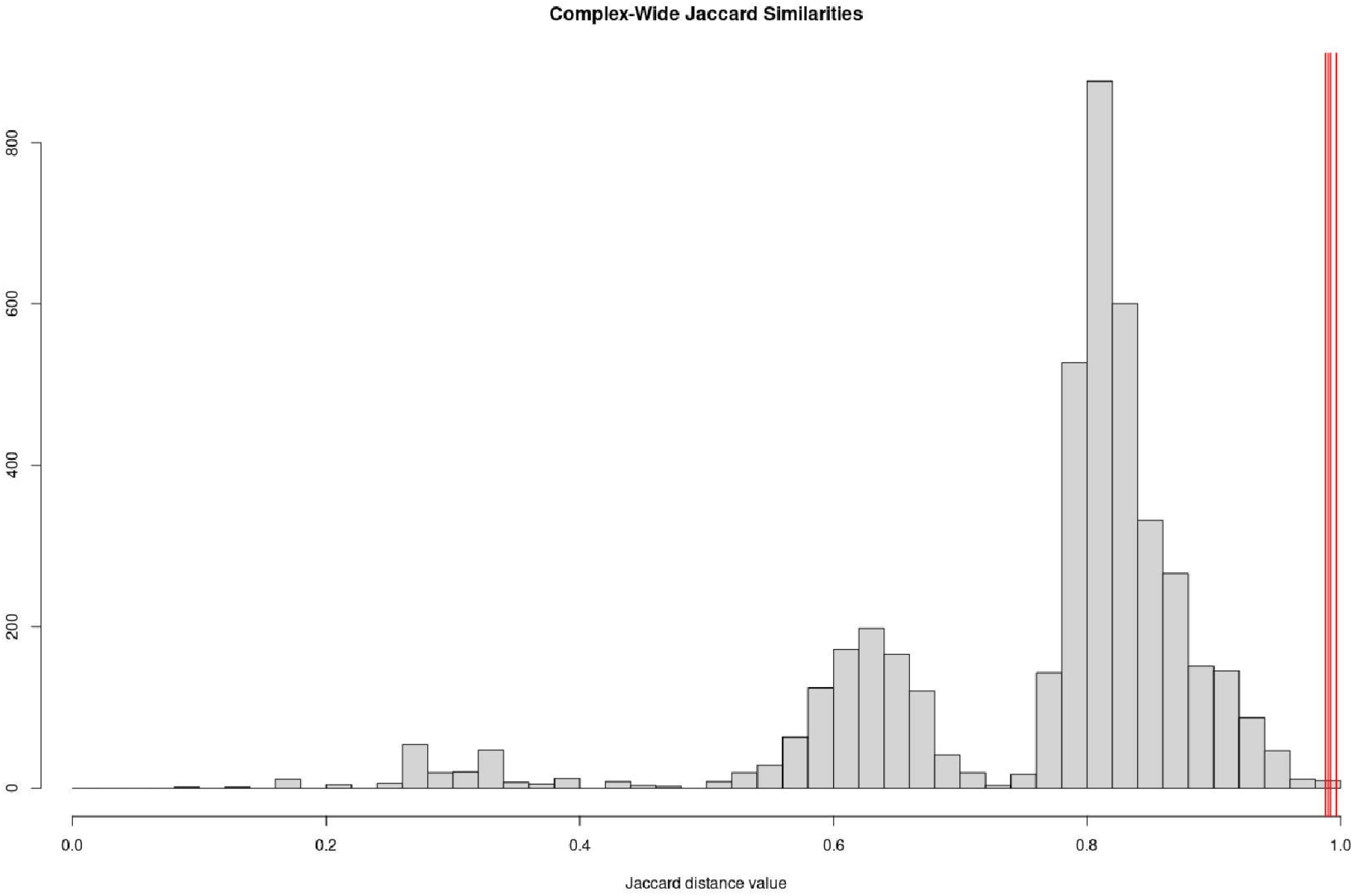
Distribution of pairwise Jaccard similarities across all samples. Similarity values of replicates are indicated by red vertical lines.

**Fig S2.**
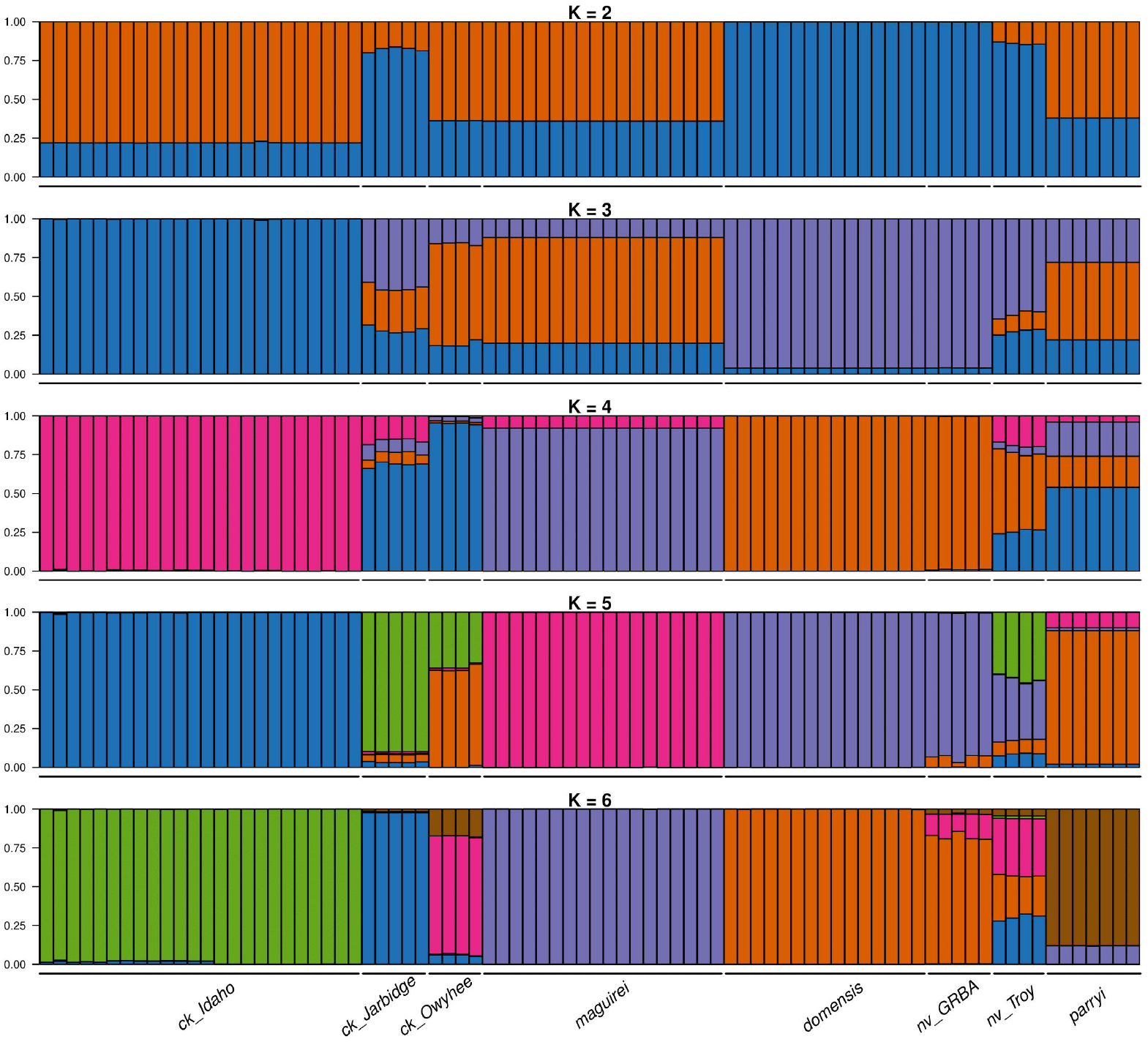
STRUCTURE plots for all samples, K values ranging from 2 to 6.

**Fig S3.**
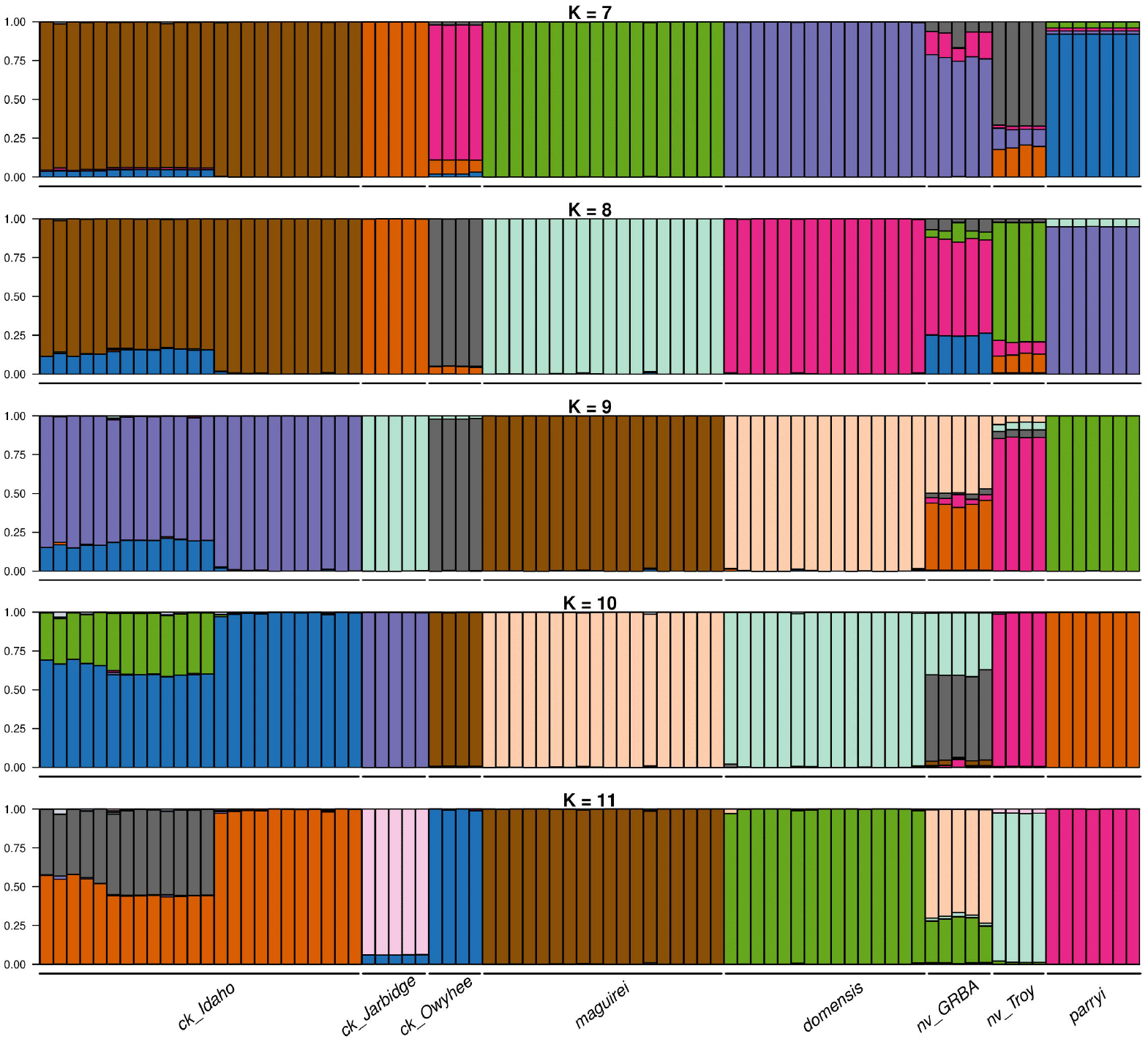
STRUCTURE plots for all samples, K values ranging from 7 to 11.

**Fig S4.**
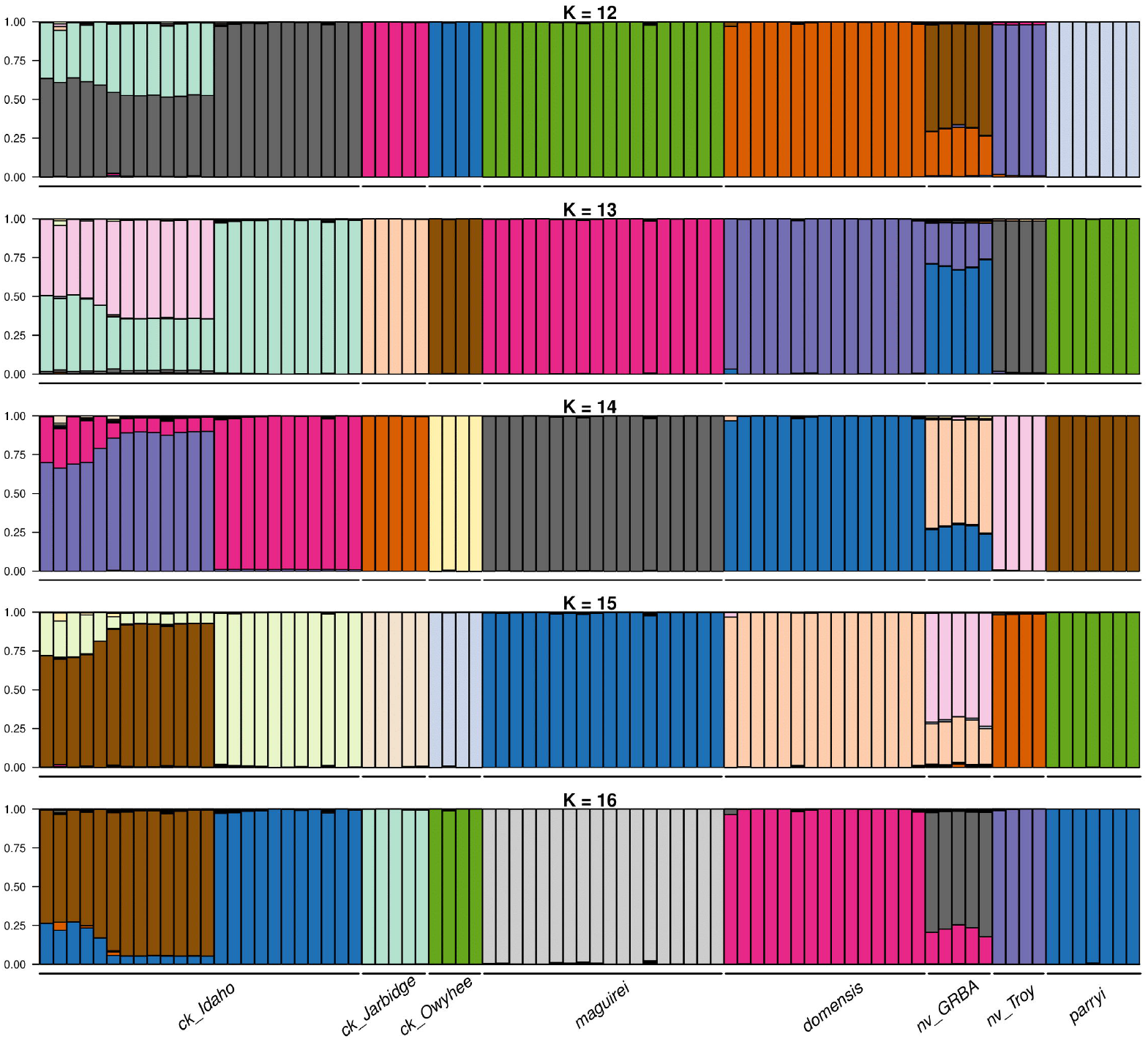
STRUCTURE plots for all samples, K values ranging from 12 to 16.

