## Supplemental Materials 1 for "Pronounced Genetic Separation Among Varieties of the *Primula cusickiana* Species Complex, a Great Basin Endemic": ChecklistProjects_ID.yml

```
http_interactions:
- request:
    method: get
    uri: http://a02235015-6.bluezone.usu.edu/api/
    body:
      encoding: ''
      string: ''
    headers:
      Accept: application/json, text/xml, application/xml, */*
  response:
    status:
      status_code: 200
      category: Success
      reason: OK
      message: 'Success: (200) OK'
    headers:
      server: nginx/1.17.4
      content-type: application/ld+json; charset=utf-8
      transfer-encoding: chunked
      connection: keep-alive
      vary: Accept
      x-content-type-options: nosniff
      x-frame-options: deny
      cache-control: no-cache, private
      date: Tue, 25 Feb 2020 17:13:00 GMT
      link: <http://a02235015-6.bluezone.usu.edu/api/docs.jsonld>; rel="http://www.w3.org/ns/hydra/core#apiDocumentation"
    body:
      encoding: UTF-8
      string: '{"@context":"\/api\/contexts\/Entrypoint","@id":"\/api","@type":"Entrypoint","configurations":"\/api\/configurations","schemaVersion":"\/api\/schemaversion","lookupLanguages":"\/api\/lookuplanguages","users":"\/api\/users","checklists":"\/api\/checklists","taxaComments":"\/api\/checklist\/taxacomments","children":"\/api\/checklist\/children","taxaLink":"\/api\/checklist\/taxalink","projectCategories":"\/api\/checklist\/projectcategories","vouchers":"\/api\/checklist\/vouchers","dynamic":"\/api\/checklist\/dynamic","checklistProjects":"\/api\/checklist\/checklistprojects","coordinates":"\/api\/checklist\/coordinates","stats":"\/api\/collection\/stats","institutions":"\/api\/collection\/institutions","collections":"\/api\/collections","categories":"\/api\/collection\/categories","queue":"\/api\/crowdsource\/queue","central":"\/api\/crowdsource\/central","titles":"\/api\/exsiccati\/titles","numbers":"\/api\/exsiccati\/numbers","sources":"\/api\/glossary\/sources","termLink":"\/api\/glossary\/termlink","glossary":"\/api\/glossary","projects":"\/api\/imageprocessor\/projects","nlpVersion":"\/api\/imageprocessor\/nlpversion","ocrFragment":"\/api\/imageprocessor\/ocrfragment","nlpFragment":"\/api\/imageprocessor\/nlpfragment","rawLabels":"\/api\/imageprocessor\/rawlabels","nlp":"\/api\/imageprocessor\/nlp","characterHeading":"\/api\/key\/characterheading","characterStateLanguages":"\/api\/key\/characterstatelanguages","characterLanguages":"\/api\/key\/characterlanguages","characters":"\/api\/key\/characters","descriptions":"\/api\/key\/descriptions","descriptionDeletions":"\/api\/key\/descriptiondeletions","characterStates":"\/api\/key\/characterstates","characterStateImages":"\/api\/key\/characterstateimages","tagKey":"\/api\/media\/tagkey","media":"\/api\/media","images":"\/api\/media\/images","guid":"\/api\/media\/guid","keywords":"\/api\/media\/keywords","tag":"\/api\/media\/tag","associations":"\/api\/occurrenceassociations\/associations","comments":"\/api\/occurrencecomments\/comments","datasetLink":"\/api\/occurrencedataset\/datasetlink","datasets":"\/api\/occurrencedataset\/datasets","genetic":"\/api\/occurrencegenetic\/genetic","exchange":"\/api\/occurrenceloan\/exchange","loans":"\/api\/occurrenceloan\/loans","revisions":"\/api\/occurrence\/revisions","edits":"\/api\/occurrence\/edits","uploadDeterminations":"\/api\/occurrence\/uploaddeterminations","fullText":"\/api\/occurrence\/fulltext","uploadParameters":"\/api\/occurrence\/uploadparameters","duplicates":"\/api\/occurrence\/duplicates","lookupChronostratigraphy":"\/api\/occurrence\/lookupchronostratigraphy","accessStats":"\/api\/occurrence\/accessstats","editLocks":"\/api\/occurrence\/editlocks","lookupCounties":"\/api\/occurrence\/lookupcounties","geoIndex":"\/api\/occurrence\/geoindex","lookupStateProvinces":"\/api\/occurrence\/lookupstateprovinces","occurrences":"\/api\/occurrences","determinations":"\/api\/occurrence\/determinations","verification":"\/api\/occurrence\/verification","uploadMappings":"\/api\/occurrence\/uploadmappings","lookupCountries":"\/api\/occurrence\/lookupcountries","identifiers":"\/api\/occurrence\/identifiers","guidOccurrences":"\/api\/occurrence\/guidoccurrences","uploadOccurrences":"\/api\/occurrence\/uploadoccurrences","guidDeterminations":"\/api\/occurrence\/guiddeterminations","lookupMunicipalities":"\/api\/occurrence\/lookupmunicipalities","checklistTaxaLink":"\/api\/reference\/checklisttaxalink","authors":"\/api\/reference\/authors","lookupReferenceTypes":"\/api\/reference\/lookupreferencetypes","references":"\/api\/references","descriptionBlock":"\/api\/taxa\/descriptionblock","synonymy":"\/api\/taxa\/synonymy","authorities":"\/api\/taxa\/authorities","taxa":"\/api\/taxa","resourceLinks":"\/api\/taxa\/resourcelinks","maps":"\/api\/taxa\/maps","descriptionStatements":"\/api\/taxa\/descriptionstatements","relationships":"\/api\/taxa\/relationships","upload":"\/api\/taxa\/upload","links":"\/api\/taxa\/links","userTaxonomy":"\/api\/taxa\/usertaxonomy","vernaculars":"\/api\/taxa\/vernaculars","ranks":"\/api\/taxa\/ranks","attributes":"\/api\/traits\/attributes","traits":"\/api\/traits","dependencies":"\/api\/traits\/dependencies","states":"\/api\/traits\/states"}'
  recorded_at: 2020-02-25 17:13:01 GMT
  recorded_with: vcr/0.3.0, webmockr/0.4.0
- request:
    method: get
    uri: http://a02235015-6.bluezone.usu.edu/api/checklist/checklistprojects/1
    body:
      encoding: ''
      string: ''
    headers:
      Accept: application/json, text/xml, application/xml, */*
  response:
    status:
      status_code: 200
      category: Success
      reason: OK
      message: 'Success: (200) OK'
    headers:
      server: nginx/1.17.4
      content-type: application/ld+json; charset=utf-8
      transfer-encoding: chunked
      connection: keep-alive
      vary: Accept
      x-content-type-options: nosniff
      x-frame-options: deny
      cache-control: private, must-revalidate
      date: Tue, 25 Feb 2020 17:13:01 GMT
      link: <http://a02235015-6.bluezone.usu.edu/api/docs.jsonld>; rel="http://www.w3.org/ns/hydra/core#apiDocumentation"
      etag: '"4766e6b77e30a36202880bc77c9027f5"'
    body:
      encoding: UTF-8
      string: '{"@context":"\/api\/contexts\/ChecklistProjects","@id":"\/api\/checklist\/checklistprojects\/1","@type":"ChecklistProjects","id":1,"projectName":"Arizona
        Flora","displayName":"Arizona Flora","managers":"Arizona State University
        Vascular Plant Herbarium","briefDescription":"Arizona is the third or fourth
        most floristically rich state in the US with perhaps as many as 3900 species
        of vascular plants. Over the last 60 years an average of ca. 12 new species
        records have been reported annually.","fullDescription":"Arizona is the third
        or fourth most floristically rich state in the US with perhaps as many as
        3900 species of vascular plants. Over the last 60 years an average of ca.
        12 new species records have been reported annually.","notes":null,"iconUrl":null,"headerUrl":null,"occurrenceSearch":0,"isPublic":1,"dynamicProperties":null,"sortSequence":10,"parentProjectId":null,"checklistId":["\/api\/checklists\/2","\/api\/checklists\/5"],"initialTimestamp":"2019-01-12T10:39:53+00:00"}'
  recorded_at: 2020-02-25 17:13:01 GMT
  recorded_with: vcr/0.3.0, webmockr/0.4.0
```
