## Supplemental Materials 1 for "Pronounced Genetic Separation Among Varieties of the *Primula cusickiana* Species Complex, a Great Basin Endemic": ChecklistProjects_page.yml

```
http_interactions:
- request:
    method: get
    uri: http://a02235015-6.bluezone.usu.edu/api/
    body:
      encoding: ''
      string: ''
    headers:
      Accept: application/json, text/xml, application/xml, */*
  response:
    status:
      status_code: 200
      category: Success
      reason: OK
      message: 'Success: (200) OK'
    headers:
      server: nginx/1.17.4
      content-type: application/ld+json; charset=utf-8
      transfer-encoding: chunked
      connection: keep-alive
      vary: Accept
      x-content-type-options: nosniff
      x-frame-options: deny
      cache-control: no-cache, private
      date: Tue, 25 Feb 2020 17:13:08 GMT
      link: <http://a02235015-6.bluezone.usu.edu/api/docs.jsonld>; rel="http://www.w3.org/ns/hydra/core#apiDocumentation"
    body:
      encoding: UTF-8
      string: '{"@context":"\/api\/contexts\/Entrypoint","@id":"\/api","@type":"Entrypoint","configurations":"\/api\/configurations","schemaVersion":"\/api\/schemaversion","lookupLanguages":"\/api\/lookuplanguages","users":"\/api\/users","checklists":"\/api\/checklists","taxaComments":"\/api\/checklist\/taxacomments","children":"\/api\/checklist\/children","taxaLink":"\/api\/checklist\/taxalink","projectCategories":"\/api\/checklist\/projectcategories","vouchers":"\/api\/checklist\/vouchers","dynamic":"\/api\/checklist\/dynamic","checklistProjects":"\/api\/checklist\/checklistprojects","coordinates":"\/api\/checklist\/coordinates","stats":"\/api\/collection\/stats","institutions":"\/api\/collection\/institutions","collections":"\/api\/collections","categories":"\/api\/collection\/categories","queue":"\/api\/crowdsource\/queue","central":"\/api\/crowdsource\/central","titles":"\/api\/exsiccati\/titles","numbers":"\/api\/exsiccati\/numbers","sources":"\/api\/glossary\/sources","termLink":"\/api\/glossary\/termlink","glossary":"\/api\/glossary","projects":"\/api\/imageprocessor\/projects","nlpVersion":"\/api\/imageprocessor\/nlpversion","ocrFragment":"\/api\/imageprocessor\/ocrfragment","nlpFragment":"\/api\/imageprocessor\/nlpfragment","rawLabels":"\/api\/imageprocessor\/rawlabels","nlp":"\/api\/imageprocessor\/nlp","characterHeading":"\/api\/key\/characterheading","characterStateLanguages":"\/api\/key\/characterstatelanguages","characterLanguages":"\/api\/key\/characterlanguages","characters":"\/api\/key\/characters","descriptions":"\/api\/key\/descriptions","descriptionDeletions":"\/api\/key\/descriptiondeletions","characterStates":"\/api\/key\/characterstates","characterStateImages":"\/api\/key\/characterstateimages","tagKey":"\/api\/media\/tagkey","media":"\/api\/media","images":"\/api\/media\/images","guid":"\/api\/media\/guid","keywords":"\/api\/media\/keywords","tag":"\/api\/media\/tag","associations":"\/api\/occurrenceassociations\/associations","comments":"\/api\/occurrencecomments\/comments","datasetLink":"\/api\/occurrencedataset\/datasetlink","datasets":"\/api\/occurrencedataset\/datasets","genetic":"\/api\/occurrencegenetic\/genetic","exchange":"\/api\/occurrenceloan\/exchange","loans":"\/api\/occurrenceloan\/loans","revisions":"\/api\/occurrence\/revisions","edits":"\/api\/occurrence\/edits","uploadDeterminations":"\/api\/occurrence\/uploaddeterminations","fullText":"\/api\/occurrence\/fulltext","uploadParameters":"\/api\/occurrence\/uploadparameters","duplicates":"\/api\/occurrence\/duplicates","lookupChronostratigraphy":"\/api\/occurrence\/lookupchronostratigraphy","accessStats":"\/api\/occurrence\/accessstats","editLocks":"\/api\/occurrence\/editlocks","lookupCounties":"\/api\/occurrence\/lookupcounties","geoIndex":"\/api\/occurrence\/geoindex","lookupStateProvinces":"\/api\/occurrence\/lookupstateprovinces","occurrences":"\/api\/occurrences","determinations":"\/api\/occurrence\/determinations","verification":"\/api\/occurrence\/verification","uploadMappings":"\/api\/occurrence\/uploadmappings","lookupCountries":"\/api\/occurrence\/lookupcountries","identifiers":"\/api\/occurrence\/identifiers","guidOccurrences":"\/api\/occurrence\/guidoccurrences","uploadOccurrences":"\/api\/occurrence\/uploadoccurrences","guidDeterminations":"\/api\/occurrence\/guiddeterminations","lookupMunicipalities":"\/api\/occurrence\/lookupmunicipalities","checklistTaxaLink":"\/api\/reference\/checklisttaxalink","authors":"\/api\/reference\/authors","lookupReferenceTypes":"\/api\/reference\/lookupreferencetypes","references":"\/api\/references","descriptionBlock":"\/api\/taxa\/descriptionblock","synonymy":"\/api\/taxa\/synonymy","authorities":"\/api\/taxa\/authorities","taxa":"\/api\/taxa","resourceLinks":"\/api\/taxa\/resourcelinks","maps":"\/api\/taxa\/maps","descriptionStatements":"\/api\/taxa\/descriptionstatements","relationships":"\/api\/taxa\/relationships","upload":"\/api\/taxa\/upload","links":"\/api\/taxa\/links","userTaxonomy":"\/api\/taxa\/usertaxonomy","vernaculars":"\/api\/taxa\/vernaculars","ranks":"\/api\/taxa\/ranks","attributes":"\/api\/traits\/attributes","traits":"\/api\/traits","dependencies":"\/api\/traits\/dependencies","states":"\/api\/traits\/states"}'
  recorded_at: 2020-02-25 17:13:09 GMT
  recorded_with: vcr/0.3.0, webmockr/0.4.0
- request:
    method: get
    uri: http://a02235015-6.bluezone.usu.edu/api/checklist/checklistprojects?page=1
    body:
      encoding: ''
      string: ''
    headers:
      Accept: application/json, text/xml, application/xml, */*
  response:
    status:
      status_code: 200
      category: Success
      reason: OK
      message: 'Success: (200) OK'
    headers:
      server: nginx/1.17.4
      content-type: application/ld+json; charset=utf-8
      transfer-encoding: chunked
      connection: keep-alive
      vary: Accept
      x-content-type-options: nosniff
      x-frame-options: deny
      cache-control: private, must-revalidate
      date: Tue, 25 Feb 2020 17:13:09 GMT
      link: <http://a02235015-6.bluezone.usu.edu/api/docs.jsonld>; rel="http://www.w3.org/ns/hydra/core#apiDocumentation"
      etag: '"9116af8271ec3e0fc507110b3cb7dddc"'
    body:
      encoding: UTF-8
      string: '{"@context":"\/api\/contexts\/ChecklistProjects","@id":"\/api\/checklist\/checklistprojects","@type":"hydra:Collection","hydra:member":[{"@id":"\/api\/checklist\/checklistprojects\/1","@type":"ChecklistProjects","id":1,"projectName":"Arizona
        Flora","displayName":"Arizona Flora","managers":"Arizona State University
        Vascular Plant Herbarium","briefDescription":"Arizona is the third or fourth
        most floristically rich state in the US with perhaps as many as 3900 species
        of vascular plants. Over the last 60 years an average of ca. 12 new species
        records have been reported annually.","fullDescription":"Arizona is the third
        or fourth most floristically rich state in the US with perhaps as many as
        3900 species of vascular plants. Over the last 60 years an average of ca.
        12 new species records have been reported annually.","notes":null,"iconUrl":null,"headerUrl":null,"occurrenceSearch":0,"isPublic":1,"dynamicProperties":null,"sortSequence":10,"parentProjectId":null,"checklistId":["\/api\/checklists\/2","\/api\/checklists\/5"],"initialTimestamp":"2019-01-12T10:39:53+00:00"},{"@id":"\/api\/checklist\/checklistprojects\/2","@type":"ChecklistProjects","id":2,"projectName":"National
        Park Service (NPS)","displayName":null,"managers":null,"briefDescription":null,"fullDescription":null,"notes":null,"iconUrl":null,"headerUrl":null,"occurrenceSearch":1,"isPublic":0,"dynamicProperties":null,"sortSequence":500,"parentProjectId":null,"checklistId":[],"initialTimestamp":"2019-01-12T10:45:17+00:00"},{"@id":"\/api\/checklist\/checklistprojects\/3","@type":"ChecklistProjects","id":3,"projectName":"National
        Park Service, Intermountain Region","displayName":"U.S. National Park","managers":"NPS","briefDescription":null,"fullDescription":"The
        collaborative on-line flora of National Park Service units of the Intermountain
        Region.","notes":null,"iconUrl":null,"headerUrl":null,"occurrenceSearch":0,"isPublic":1,"dynamicProperties":null,"sortSequence":54,"parentProjectId":"\/api\/checklist\/checklistprojects\/2","checklistId":["\/api\/checklists\/2","\/api\/checklists\/6"],"initialTimestamp":"2019-01-12T10:39:53+00:00"},{"@id":"\/api\/checklist\/checklistprojects\/4","@type":"ChecklistProjects","id":4,"projectName":"Flora
        of the National Park Service - Southwestern US","displayName":null,"managers":"Steve
        Buckley, SODN-NPS","briefDescription":null,"fullDescription":null,"notes":null,"iconUrl":null,"headerUrl":null,"occurrenceSearch":0,"isPublic":0,"dynamicProperties":null,"sortSequence":110,"parentProjectId":"\/api\/checklist\/checklistprojects\/2","checklistId":["\/api\/checklists\/2"],"initialTimestamp":"2019-01-12T10:42:31+00:00"},{"@id":"\/api\/checklist\/checklistprojects\/5","@type":"ChecklistProjects","id":5,"projectName":"Colorado
        Plateau","displayName":null,"managers":"Tina Ayers, Susan Holiday, Glenn Rink","briefDescription":null,"fullDescription":null,"notes":null,"iconUrl":null,"headerUrl":null,"occurrenceSearch":0,"isPublic":1,"dynamicProperties":null,"sortSequence":48,"parentProjectId":null,"checklistId":["\/api\/checklists\/5"],"initialTimestamp":"2019-01-12T10:43:33+00:00"},{"@id":"\/api\/checklist\/checklistprojects\/6","@type":"ChecklistProjects","id":6,"projectName":"Tucson
        Flora","displayName":null,"managers":"Tucson SEINet Community Effort","briefDescription":null,"fullDescription":"This
        project is being developed as a community volunteer project. By periodically
        meeting in small groups, we have started to systematically inventory public
        places located within the Tucson Region. If you wish to participate or lead
        a group, for more information.","notes":null,"iconUrl":null,"headerUrl":null,"occurrenceSearch":0,"isPublic":0,"dynamicProperties":null,"sortSequence":50,"parentProjectId":null,"checklistId":["\/api\/checklists\/3"],"initialTimestamp":"2019-01-12T10:44:39+00:00"},{"@id":"\/api\/checklist\/checklistprojects\/7","@type":"ChecklistProjects","id":7,"projectName":"Teaching
        Checklists - Western Regions","displayName":null,"managers":"Misc Managers","briefDescription":null,"fullDescription":null,"notes":null,"iconUrl":null,"headerUrl":null,"occurrenceSearch":0,"isPublic":1,"dynamicProperties":null,"sortSequence":60,"parentProjectId":null,"checklistId":["\/api\/checklists\/12"],"initialTimestamp":"2019-01-12T10:39:53+00:00"},{"@id":"\/api\/checklist\/checklistprojects\/8","@type":"ChecklistProjects","id":8,"projectName":"Bureau
        of Land Management (BLM)","displayName":null,"managers":null,"briefDescription":null,"fullDescription":null,"notes":null,"iconUrl":null,"headerUrl":null,"occurrenceSearch":1,"isPublic":0,"dynamicProperties":null,"sortSequence":510,"parentProjectId":null,"checklistId":[],"initialTimestamp":"2019-01-12T10:45:53+00:00"},{"@id":"\/api\/checklist\/checklistprojects\/9","@type":"ChecklistProjects","id":9,"projectName":"United
        States Forest Service (USFS)","displayName":null,"managers":null,"briefDescription":null,"fullDescription":null,"notes":null,"iconUrl":null,"headerUrl":null,"occurrenceSearch":1,"isPublic":0,"dynamicProperties":null,"sortSequence":520,"parentProjectId":null,"checklistId":[],"initialTimestamp":"2019-01-12T10:46:31+00:00"},{"@id":"\/api\/checklist\/checklistprojects\/10","@type":"ChecklistProjects","id":10,"projectName":"Chicago
        Region Checklists and Inventories","displayName":null,"managers":"Andrew Hipp","briefDescription":null,"fullDescription":null,"notes":null,"iconUrl":null,"headerUrl":null,"occurrenceSearch":0,"isPublic":1,"dynamicProperties":null,"sortSequence":40,"parentProjectId":null,"checklistId":["\/api\/checklists\/10"],"initialTimestamp":"2019-01-12T10:46:51+00:00"},{"@id":"\/api\/checklist\/checklistprojects\/11","@type":"ChecklistProjects","id":11,"projectName":"Macrofungi
        of North America: Regions","displayName":null,"managers":null,"briefDescription":null,"fullDescription":"\u003Cp
        align=\u0027justify\u0027\u003E As perhaps only 5% of the total diversity
        of fungal species on Earth have been described (Hawksworth D.L. \u003Ci\u003EMycological
        Research\u003C\/i\u003E 105:1422-1432), the checklists presented here are
        likely far from being complete. These checklists, therefore, are works in
        progress and should be considered contributions toward a more comprehensive
        accounting of the diversity of macrofungal species within North America.\u003C\/p\u003E","notes":null,"iconUrl":null,"headerUrl":null,"occurrenceSearch":0,"isPublic":1,"dynamicProperties":null,"sortSequence":6,"parentProjectId":null,"checklistId":["\/api\/checklists\/14","\/api\/checklists\/17"],"initialTimestamp":"2019-01-12T17:03:50+00:00"},{"@id":"\/api\/checklist\/checklistprojects\/12","@type":"ChecklistProjects","id":12,"projectName":"Macrofungi
        of North America: Groups","displayName":null,"managers":null,"briefDescription":null,"fullDescription":"\u003Cp
        align=\u0027justify\u0027\u003EAs perhaps only 5% of the total diversity of
        fungal species on Earth have been described (Hawksworth D.L. \u003Ci\u003EMycological
        Research\u003C\/i\u003E 105:1422-1432), the checklists presented here are
        likely far from being complete. These checklists, therefore, are works in
        progress, and should be considered contributions toward a more comprehensive
        accounting of the diversity of macrofungal species within North America.\u003C\/p\u003E","notes":null,"iconUrl":null,"headerUrl":null,"occurrenceSearch":0,"isPublic":1,"dynamicProperties":null,"sortSequence":4,"parentProjectId":null,"checklistId":[],"initialTimestamp":"2019-01-12T17:03:50+00:00"},{"@id":"\/api\/checklist\/checklistprojects\/13","@type":"ChecklistProjects","id":13,"projectName":"Fungi
        of North America: Historic","displayName":null,"managers":null,"briefDescription":null,"fullDescription":"\u003Cp
        align=\u0027justify\u0027\u003EAs perhaps only 5% of the total diversity of
        fungal species on Earth have been described (Hawksworth D.L. \u003Ci\u003EMycological
        Research\u003C\/i\u003E 105:1422-1432), knowledge of the fungal flora of the
        United States is likely far from being complete. The checklists presented
        here represent historic published accounts or list of species authored by
        mycologists who described large numbers of taxa from North America. Acknowledged
        here are the considerable contributions that the mycologists associated with
        these lists made toward a more comprehensive accounting of the diversity of
        macrofungal species within North America.\u003C\/p\u003E","notes":null,"iconUrl":null,"headerUrl":null,"occurrenceSearch":0,"isPublic":1,"dynamicProperties":null,"sortSequence":2,"parentProjectId":null,"checklistId":["\/api\/checklists\/15","\/api\/checklists\/16"],"initialTimestamp":"2019-01-12T17:03:50+00:00"},{"@id":"\/api\/checklist\/checklistprojects\/14","@type":"ChecklistProjects","id":14,"projectName":"Macrofungi
        Across the World: Countries","displayName":null,"managers":null,"briefDescription":null,"fullDescription":"\u003Cp
        align=\u0027justify\u0027\u003E As perhaps only 5% of the total diversity
        of fungal species on Earth have been described (Hawksworth D.L. Mycological
        Research 105:1422-1432), the checklists presented here are likely far from
        being complete. These checklists, therefore, are works in progress and should
        be considered contributions toward a more comprehensive accounting of the
        worldwide diversity of macrofungal species.\u003C\/p\u003E","notes":null,"iconUrl":null,"headerUrl":null,"occurrenceSearch":0,"isPublic":1,"dynamicProperties":null,"sortSequence":3,"parentProjectId":null,"checklistId":[],"initialTimestamp":"2019-01-12T17:03:50+00:00"},{"@id":"\/api\/checklist\/checklistprojects\/15","@type":"ChecklistProjects","id":15,"projectName":"Macrofungi
        of North America: Local","displayName":null,"managers":null,"briefDescription":null,"fullDescription":"\u003Cp
        align=\u0027justify\u0027\u003EAs perhaps only 5% of the total diversity of
        fungal species on Earth have been described (Hawksworth D.L. \u003Ci\u003EMycological
        Research\u003C\/i\u003E 105:1422-1432), the checklists presented here are
        likely far from being complete. These checklists, therefore, are works in
        progress and should be considered contributions toward a more comprehensive
        accounting of the diversity of macrofungal species within North America.\u003C\/p\u003E","notes":null,"iconUrl":null,"headerUrl":null,"occurrenceSearch":0,"isPublic":1,"dynamicProperties":null,"sortSequence":5,"parentProjectId":null,"checklistId":[],"initialTimestamp":"2019-01-12T17:03:50+00:00"},{"@id":"\/api\/checklist\/checklistprojects\/16","@type":"ChecklistProjects","id":16,"projectName":"Microfungi
        of North America: Regions","displayName":null,"managers":null,"briefDescription":null,"fullDescription":"As
        perhaps only 5% of the total diversity of fungal species on Earth have been
        described (Hawksworth D.L. Mycological Research 105:1422-1432), the checklists
        presented here are likely far from being complete. These checklists, therefore,
        are works in progress and should be considered contributions toward a more
        comprehensive accounting of the diversity of macrofungal species within North
        America.","notes":null,"iconUrl":null,"headerUrl":null,"occurrenceSearch":0,"isPublic":1,"dynamicProperties":null,"sortSequence":8,"parentProjectId":null,"checklistId":[],"initialTimestamp":"2019-01-12T17:03:50+00:00"},{"@id":"\/api\/checklist\/checklistprojects\/17","@type":"ChecklistProjects","id":17,"projectName":"Microfungi
        Across the World: Countries","displayName":null,"managers":null,"briefDescription":null,"fullDescription":"\u003Cp
        align=\u0027justify\u0027\u003E As perhaps only 5% of the total diversity
        of fungal species on Earth have been described (Hawksworth D.L. Mycological
        Research 105:1422-1432), the checklists presented here are likely far from
        being complete. These checklists, therefore, are works in progress and should
        be considered contributions toward a more comprehensive accounting of the
        worldwide diversity of microfungal species.\u003C\/p\u003E","notes":null,"iconUrl":null,"headerUrl":null,"occurrenceSearch":0,"isPublic":1,"dynamicProperties":null,"sortSequence":7,"parentProjectId":null,"checklistId":[],"initialTimestamp":"2019-01-12T17:03:50+00:00"},{"@id":"\/api\/checklist\/checklistprojects\/18","@type":"ChecklistProjects","id":18,"projectName":"Fungi
        Across the World: Historic","displayName":null,"managers":null,"briefDescription":null,"fullDescription":"\u003Cp
        align=\u0027justify\u0027\u003EAs perhaps only 5% of the total diversity of
        fungal species on Earth have been described (Hawksworth D.L. \u003Ci\u003EMycological
        Research\u003C\/i\u003E 105:1422-1432), knowledge of the fungal flora of the
        Earth is likely far from being complete. The checklists presented here represent
        historic published accounts or list of species authored by mycologists who
        described large numbers of taxa or are known for collecting in a particular
        regions of the World. Acknowledged here are the considerable contributions
        that the mycologists associated with these lists made toward a more comprehensive
        accounting of the diversity of macrofungal species across the globe.\u003C\/p\u003E","notes":null,"iconUrl":null,"headerUrl":null,"occurrenceSearch":0,"isPublic":1,"dynamicProperties":null,"sortSequence":1,"parentProjectId":null,"checklistId":[],"initialTimestamp":"2019-01-12T17:03:50+00:00"}],"hydra:totalItems":18}'
  recorded_at: 2020-02-25 17:13:09 GMT
  recorded_with: vcr/0.3.0, webmockr/0.4.0
```
