## Supplemental Materials 1 for "Pronounced Genetic Separation Among Varieties of the *Primula cusickiana* Species Complex, a Great Basin Endemic": README.md

# vcf2Jaccard
Converts the variant-calling files (vcf) from ipyrad output to a sample pairwise dataframe of mean Jaccard similarity coefficients (aka Jaccard index, or Intersection over Union) across all SNPs.
