## Supplemental Materials 1 for "Pronounced Genetic Separation Among Varieties of the *Primula cusickiana* Species Complex, a Great Basin Endemic": vcf2Jaccard.html


### vcf2Jaccard

#### vcf2Jaccard.py

vcf2Jaccard2.py

```
"""
------------------------------------------------------------------------------------------
Description: Converts the variant-calling files (vcf) from ipyrad output to a dataframe of mean Jaccard similarity coefficients (aka Jaccard index, or Intersection over Union) across all SNPs.

Usage: python vcf2Jaccard.py your_input.vcf
2 optional arguements: -N, -o. For more information: see bottom

File Name: vcf2jacsim.py
Author: Carol Rowe
Date Created: 2019-05-3 using Python 3.7.1


NOTES:
For all sample pairwise comparisons:
Jaccard similarity coefficients are calculated for each SNP.
A mean is then taken across all the SNPs, with missing data ignored.
This mean is then entered into the final similarity matrix (dataframe).

Three output files are created:
1) Jaccob_sim_means.csv
    You have the option to change this file name when you run the script.
    Pairwise similarity matrix with mean Jaccard similarity coefficients
2) SNP_tally.csv
    Pairwise matrix where values are counts of non-missing SNPs
3) missing_SNP.csv
    Pairwise matrix where values are counts of missing SNPs
-----------------------------------------------------------------------------------------
"""

import argparse
import pandas as pd
from itertools import combinations
import numpy as np
import copy

__author__ = "Carol Rowe"


def vcf2Jaccard(filename, skiplines, outfile):  
    # DF = DATAFRAME
    # READ IN THE FILE AS A DF AND GET JUST THE NEEDED COLUMNS
    # read in the .vcf file from the ipyrad output into a df (dataframe)
    vcf = pd.read_csv(filename, sep="\t", skiprows=skiplines)
    # Slice df: keep just columns with sample data
    data = vcf.iloc[0:,9:]
    # The data contains "." which is missing data. These periods will be problematic for downstram steps.
    # Using regular expressions to remove the ".".
    # Three scenarios: "./.", "0/.", or "./0" (The 0 can be any other integer [0-4])
    # Replacing the "." with a 9 to keep all as integers.
    # Remember, 9 now represents missing data.
    data = data.replace({"\./\." : "9/9"}, regex=True)
    data= data.replace({"[0-9]/(\.)" : "9/0"}, regex=True)
    data = data.replace({"(\.)/[0-9]" : "0/9"}, regex=True)
    # Each cell now has format: 1/1:12:11,0,1,0
    # Separated by ":", we have allele ref, depth coverage, and counts for each AGCT
    # Need just allele ref. (i.e. is it homozygous, hetero, and for which dNTP)
    format = lambda x: x.split(':')[0]
    data2 = data.applymap(format)
    # Want that into a list. Now each cell in df will have something like: [0, 0]
    format = lambda x: x.split('/')
    data3 = data2.applymap(format)

    # python has a library for jaccard similarity, but I need to be able to handle missing data.
    # Remember, our missing data is now: 9
    def jaccard_similarity(list1, list2):
        if ("9" in list1) or ("9" in list2):
            jac_sim = "NA"
        else:
            inter = len(set(list1).intersection(set(list2)))
            union = len(set(list1).union(set(list2)))
            jac_sim = inter/union
        return jac_sim

    # When we use the Jaccard similiarity function, we will also be generating a list.
    # This list will have missing values now as "NA" 
    # We do not want to use the missing values when calculating means.
    def remove_NA_from_list(list):
        return [e for e in list if e != 'NA']

    # GET SAMPLE NAMES TO A LIST
    # get the column names (your sample names) as a list
    sample_names = data3.columns.values.tolist()
  
    # use itertools.combinations to get all pairwise combos of samples without being repetitive.
    # in other words, we want sample A to B combo, but don't need B to A, or A to A, or B to B.
    sample_combo = list(combinations(sample_names,2))

    # Create a new, empty df which we will be adding the Jaccob similairty means to
    jac_sim_df = pd.DataFrame(index=sample_names, columns=sample_names)
    tally_df = pd.DataFrame(index=sample_names, columns=sample_names)
    missing_df = pd.DataFrame(index=sample_names, columns=sample_names)

    for compare in sample_combo:
        col1 = compare[0]
        col2 = compare[1]
        list1 = data3[col1].values.tolist()
        list2 = data3[col2].values.tolist()
        js_list = []
        for i in range(0,len(list1)):
            value = jaccard_similarity(list1[i], list2[i])
            js_list.append(value)
        new_list = remove_NA_from_list(js_list)
        # If we end up with an empty list, make sure the mean is "NA"
        if len(new_list) == 0:
            js_mean = "NA"
        # Get the mean of the non-NA values in the list
        else:
            js_mean = np.mean(new_list)
        # Enter the js_mean score to the correct location in the pairwise dataframe
        jac_sim_df.loc[col1,col2] = js_mean
        # Enter the total number of non-NA pairwise SNPs into the SNP_tally matrix
        tally_df.loc[col1,col2] = len(new_list)
        # Calculate the number of NA SNPs and enter into the misssing_SNP datadframe
        missing = len(js_list) - len(new_list)
        missing_df.loc[col1,col2] = missing

    # Save your final dfs to files:
    jac_sim_df.to_csv(outfile)
    tally_df.to_csv("SNP_tally.csv")
    missing_df.to_csv("missing_SNP.csv")

    input_Count_Row=data3.shape[0] #gives number of row count
    input_Count_Col=data3.shape[1] #gives number of col count
    print("Your input vcf data file contains {} samples and {} SNPs." .format(input_Count_Col, input_Count_Row))
    print("Your Jaccard similarity coefficient means have now been caluculated.")


# if name in main so we can run vcf2Jacsim.py by itself (main)
# or it can still be used if you use it embedded (import vcf2Jacsim) within another script (name)
if __name__ in '__main__':
    # This allows the --help to show the docstring at the top of this script.
    parser = argparse.ArgumentParser(description=__doc__, formatter_class=argparse.RawDescriptionHelpFormatter)
    # Add arguments (3):
    # First argument is mandatory
    parser.add_argument('input', metavar='vcf_input_file', help='Enter the name of your vcf file.')
    # Next 2 arguments are optional
    parser.add_argument('-N','--skiplines', help='Number of lines to skip in vcf file until header of the data table. Default = 10', type=int, default=10, required=False)
    parser.add_argument('-o','--outfile', help='Output Jaccard similarity means dataframe. Default = Jaccob_sim_means.csv', type=str, default='Jaccob_sim_means.csv', required=False)
    # Array for all arguments passed to script:
    args = parser.parse_args()

    # Now, we can access the arguments input by the user (or use defaults), and apply to our function   
    vcf2Jaccard(args.input, args.skiplines, args.outfile)
```

#### Extra details of vcf2Jaccard.py

This file is just for additional output of some intermediate steps of the vcf2Jaccard.py script.

Reminder on how to run:  
python vcf2Jaccard.py your\_file.vcf  
For help menu:  
python vcf2Jaccard.py -h

```
# created a reduced size .vcf file to play with: temp2.vcf 
# read in file
vcf = pd.read_csv("temp2.vcf", sep="\t", skiprows=10)
# Slice df: keep just columns with sample data
data = vcf.iloc[0:,9:]
print(vcf.iloc[0:7,0:12])
'''
        ABEY-034-18      ABPZ-076-18      ABVE-221-18      ACFL-031-18  \
0   0/0:10:0,0,10,0  0/1:13:3,0,10,0  0/0:13:0,0,13,0  0/1:17:4,0,13,0   
1   0/0:10:10,0,0,0  0/0:15:15,0,0,0  0/0:13:13,0,0,0  0/0:16:15,0,1,0   
2   0/0:10:0,0,10,0  0/0:15:0,0,15,0  0/0:13:0,0,13,0  0/0:17:0,0,17,0   
3   0/0:10:0,10,0,0  0/0:15:0,15,0,0  0/0:13:0,13,0,0  0/0:17:0,17,0,0   
4   0/0:10:0,0,10,0  0/0:15:0,0,15,0  0/0:13:0,0,13,0  0/0:17:0,0,17,0   
5   0/0:10:10,0,0,0  0/0:15:15,0,0,0  0/0:13:13,0,0,0  0/0:17:17,0,0,0   
6  1/0:35:18,0,17,0  0/0:48:48,0,0,0  0/0:11:11,0,0,0  1/0:22:13,0,9,0   

    ACGNT01-497-18   AEBVG-419-18      AFOG-113-18     AGXLZ-327-18  \
0  0/0:10:0,0,10,0  ./.:0:0,0,0,0  1/1:12:11,0,1,0  0/0:17:0,0,17,0   
1  0/0:10:10,0,0,0  ./.:0:0,0,0,0  0/0:12:12,0,0,0  1/0:17:12,0,5,0   
2  0/0:10:0,0,10,0  ./.:0:0,0,0,0  0/0:12:0,0,12,0  0/0:17:0,0,17,0   
3  0/0:10:0,10,0,0  ./.:0:0,0,0,0  0/0:12:1,11,0,0  0/0:17:0,17,0,0   
4  0/0:10:0,0,10,0  ./.:0:0,0,0,0  0/0:12:0,0,12,0  1/0:17:0,0,12,5   
5  0/0:10:10,0,0,0  ./.:0:0,0,0,0  0/0:12:12,0,0,0  0/0:17:17,0,0,0   
6  1/1:18:0,0,18,0  0/0:9:9,0,0,0  0/0:26:26,0,0,0  0/0:27:26,0,1,0   

       AHQT-229-18    AHUK-131-18      AHYY-234-18     AMYCG-281-18  
0  ./.:18:3,0,15,0  0/0:7:0,0,7,0  0/0:27:0,0,27,0    ./.:0:0,0,0,0  
1  0/0:18:18,0,0,0  0/0:7:7,0,0,0  0/0:27:27,0,0,0    ./.:0:0,0,0,0  
2  0/0:18:0,0,18,0  0/0:7:0,0,7,0  0/0:27:0,0,27,0    ./.:0:0,0,0,0  
3  0/0:18:0,18,0,0  0/0:7:0,7,0,0  0/0:27:0,27,0,0    ./.:0:0,0,0,0  
4  0/0:18:0,0,18,0  0/0:7:0,0,7,0  0/0:27:0,0,27,0    ./.:0:0,0,0,0  
5  0/0:18:18,0,0,0  0/0:7:7,0,0,0  0/0:27:27,0,0,0    ./.:0:0,0,0,0  
6  0/0:36:36,0,0,0  0/0:9:9,0,0,0  0/0:28:28,0,0,0  0/0:26:26,0,0,0  
'''

# The data contains "." which is missing data. These periods will be problematic for downstram steps.
# Using regular expressions to remove the ".".
# Three scenarios: "./.", "0/.", or "./0" (The 0 can be any other integer [0-4])
# Replacing the "." with a 9 to keep all as integers.
# Remember, 9 now represents missing data.
data = data.replace({"\./\." : "9/9"}, regex=True)
data= data.replace({"[0-9]/(\.)" : "9/0"}, regex=True)
data = data.replace({"(\.)/[0-9]" : "0/9"}, regex=True)
print(data.iloc[0:7,0:7])
'''
        ABEY-034-18      ABPZ-076-18      ABVE-221-18      ACFL-031-18  \
0   0/0:10:0,0,10,0  0/1:13:3,0,10,0  0/0:13:0,0,13,0  0/1:17:4,0,13,0   
1   0/0:10:10,0,0,0  0/0:15:15,0,0,0  0/0:13:13,0,0,0  0/0:16:15,0,1,0   
2   0/0:10:0,0,10,0  0/0:15:0,0,15,0  0/0:13:0,0,13,0  0/0:17:0,0,17,0   
3   0/0:10:0,10,0,0  0/0:15:0,15,0,0  0/0:13:0,13,0,0  0/0:17:0,17,0,0   
4   0/0:10:0,0,10,0  0/0:15:0,0,15,0  0/0:13:0,0,13,0  0/0:17:0,0,17,0   
5   0/0:10:10,0,0,0  0/0:15:15,0,0,0  0/0:13:13,0,0,0  0/0:17:17,0,0,0   
6  1/0:35:18,0,17,0  0/0:48:48,0,0,0  0/0:11:11,0,0,0  1/0:22:13,0,9,0   

    ACGNT01-497-18   AEBVG-419-18      AFOG-113-18  
0  0/0:10:0,0,10,0  9/9:0:0,0,0,0  1/1:12:11,0,1,0  
1  0/0:10:10,0,0,0  9/9:0:0,0,0,0  0/0:12:12,0,0,0  
2  0/0:10:0,0,10,0  9/9:0:0,0,0,0  0/0:12:0,0,12,0  
3  0/0:10:0,10,0,0  9/9:0:0,0,0,0  0/0:12:1,11,0,0  
4  0/0:10:0,0,10,0  9/9:0:0,0,0,0  0/0:12:0,0,12,0  
5  0/0:10:10,0,0,0  9/9:0:0,0,0,0  0/0:12:12,0,0,0  
6  1/1:18:0,0,18,0  0/0:9:9,0,0,0  0/0:26:26,0,0,0  
'''

# Each cell now has format: 1/1:12:11,0,1,0
# Separated by ":", we have allele ref, depth coverage, and counts for each AGCT
# Need just allele ref. (i.e. is it homozygous, hetero, and for which dNTP)
format = lambda x: x.split(':')[0]
data2 = data.applymap(format)
print(data2.iloc[0:7,0:7])
'''
 ABEY-034-18 ABPZ-076-18 ABVE-221-18 ACFL-031-18 ACGNT01-497-18 AEBVG-419-18  \
0         0/0         0/1         0/0         0/1            0/0          9/9   
1         0/0         0/0         0/0         0/0            0/0          9/9   
2         0/0         0/0         0/0         0/0            0/0          9/9   
3         0/0         0/0         0/0         0/0            0/0          9/9   
4         0/0         0/0         0/0         0/0            0/0          9/9   
5         0/0         0/0         0/0         0/0            0/0          9/9   
6         1/0         0/0         0/0         1/0            1/1          0/0   

  AFOG-113-18  
0         1/1  
1         0/0  
2         0/0  
3         0/0  
4         0/0  
5         0/0  
6         0/0 
'''

# Want that into a list. Now each cell in df will have something like: [0, 0]
format = lambda x: x.split('/')
data3 = data2.applymap(format)
print(data3.iloc[0:7,0:7])
'''
  ABEY-034-18 ABPZ-076-18 ABVE-221-18 ACFL-031-18 ACGNT01-497-18 AEBVG-419-18  \
0      [0, 0]      [0, 1]      [0, 0]      [0, 1]         [0, 0]       [9, 9]   
1      [0, 0]      [0, 0]      [0, 0]      [0, 0]         [0, 0]       [9, 9]   
2      [0, 0]      [0, 0]      [0, 0]      [0, 0]         [0, 0]       [9, 9]   
3      [0, 0]      [0, 0]      [0, 0]      [0, 0]         [0, 0]       [9, 9]   
4      [0, 0]      [0, 0]      [0, 0]      [0, 0]         [0, 0]       [9, 9]   
5      [0, 0]      [0, 0]      [0, 0]      [0, 0]         [0, 0]       [9, 9]   
6      [1, 0]      [0, 0]      [0, 0]      [1, 0]         [1, 1]       [0, 0]   

  AFOG-113-18  
0      [1, 1]  
1      [0, 0]  
2      [0, 0]  
3      [0, 0]  
4      [0, 0]  
5      [0, 0]  
6      [0, 0]  
'''

# python has a library for jaccard similarity, but I need to be able to handle missing data.
# Remember, our missing data is now: 9
def jaccard_similarity(list1, list2):
    if ("9" in list1) or ("9" in list2):
        jac_sim = "NA"
    else:
        inter = len(set(list1).intersection(set(list2)))
        union = len(set(list1).union(set(list2)))
        jac_sim = inter/union
    return jac_sim

# When we use the Jaccard similiarity function, we will also be generating a list.
# This list will have missing values now as "NA" 
# We do not want to use the missing values when calculating means.
def remove_NA_from_list(list):
    return [e for e in list if e != 'NA']

# GET SAMPLE NAMES TO A LIST
# get the column names (your sample names) as a list
sample_names = data3.columns.values.tolist()
print(len(sample_names))
print(sample_names[0:6])
'''
536
['ABEY-034-18', 'ABPZ-076-18', 'ABVE-221-18', 'ACFL-031-18', 'ACGNT01-497-18', 'AEBVG-419-18']
'''

# use itertools.combinations to get all pairwise combos of samples without being repetitive.
# in other words, we want sample A to B combo, but don't need B to A, or A to A, or B to B.
sample_combo = list(combinations(sample_names,2))
print(len(sample_combo))
print(sample_combo[0:6])
'''
143380
[('ABEY-034-18', 'ABPZ-076-18'), ('ABEY-034-18', 'ABVE-221-18'), ('ABEY-034-18', 'ACFL-031-18'), ('ABEY-034-18', 'ACGNT01-497-18'), ('ABEY-034-18', 'AEBVG-419-18'), ('ABEY-034-18', 'AFOG-113-18')]
'''

# Create a new, empty df which we will be adding the Jaccob similairty means to
jac_sim_df = pd.DataFrame(index=sample_names, columns=sample_names)
print(jac_sim_df.iloc[0:7,0:7])
'''
               ABEY-034-18 ABPZ-076-18 ABVE-221-18 ACFL-031-18 ACGNT01-497-18  \
ABEY-034-18            NaN         NaN         NaN         NaN            NaN   
ABPZ-076-18            NaN         NaN         NaN         NaN            NaN   
ABVE-221-18            NaN         NaN         NaN         NaN            NaN   
ACFL-031-18            NaN         NaN         NaN         NaN            NaN   
ACGNT01-497-18         NaN         NaN         NaN         NaN            NaN   
AEBVG-419-18           NaN         NaN         NaN         NaN            NaN   
AFOG-113-18            NaN         NaN         NaN         NaN            NaN   

               AEBVG-419-18 AFOG-113-18  
ABEY-034-18             NaN         NaN  
ABPZ-076-18             NaN         NaN  
ABVE-221-18             NaN         NaN  
ACFL-031-18             NaN         NaN  
ACGNT01-497-18          NaN         NaN  
AEBVG-419-18            NaN         NaN  
AFOG-113-18             NaN         NaN  
'''
# These two empty dataframes look just as the example above
tally_df = pd.DataFrame(index=sample_names, columns=sample_names)
missing_df = pd.DataFrame(index=sample_names, columns=sample_names)


for compare in sample_combo:
    col1 = compare[0]
    col2 = compare[1]
    list1 = data3[col1].values.tolist()
    list2 = data3[col2].values.tolist()
    js_list = []
    for i in range(0,len(list1)):
        value = jaccard_similarity(list1[i], list2[i])
        js_list.append(value)
    new_list = remove_NA_from_list(js_list)
    # If we end up with an empty list, make sure the mean is "NA"
    if len(new_list) == 0:
        js_mean = "NA"
    # Get the mean of the non-NA values in the list
    else:
        js_mean = np.mean(new_list)
    # Enter the js_mean score to the correct location in the pairwise dataframe
    jac_sim_df.loc[col1,col2] = js_mean
    # Enter the total number of non-NA pairwise SNPs into the SNP_tally matrix
    tally_df.loc[col1,col2] = len(new_list)
    # Calculate the number of NA SNPs and enter into the misssing_SNP datadframe
    missing = len(js_list) - len(new_list)
    missing_df.loc[col1,col2] = missing

# See what the output dataframes look like
print(jac_sim_df.iloc[0:7,0:10])
'''
               ABEY-034-18 ABPZ-076-18 ABVE-221-18 ACFL-031-18 ACGNT01-497-18  \
ABEY-034-18            NaN    0.921053    0.944444    0.973684       0.842105   
ABPZ-076-18            NaN         NaN    0.944444    0.947368       0.815789   
ABVE-221-18            NaN         NaN         NaN    0.888889       0.833333   
ACFL-031-18            NaN         NaN         NaN         NaN       0.815789   
ACGNT01-497-18         NaN         NaN         NaN         NaN            NaN   
AEBVG-419-18           NaN         NaN         NaN         NaN            NaN   
AFOG-113-18            NaN         NaN         NaN         NaN            NaN   

               AEBVG-419-18 AFOG-113-18 AGXLZ-327-18 AHQT-229-18 AHUK-131-18  
ABEY-034-18        0.833333    0.894737     0.894737    0.941176    0.947368  
ABPZ-076-18               1    0.973684     0.921053           1    0.973684  
ABVE-221-18               1    0.888889     0.888889           1           1  
ACFL-031-18        0.833333    0.921053     0.868421    0.941176    0.921053  
ACGNT01-497-18          0.5    0.789474     0.789474    0.852941    0.842105  
AEBVG-419-18            NaN           1            1           1           1  
AFOG-113-18             NaN         NaN     0.894737           1    0.947368  
'''

# Save your final df to a file:
jac_sim_df.to_csv(outfile)
tally_df.to_csv("SNP_tally.csv")
missing_df.to_csv("missing_SNP.csv")

# Print general info for user
input_Count_Row=data3.shape[0] #gives number of row count
input_Count_Col=data3.shape[1] #gives number of col count
print("Your input vcf data file contains {} samples and {} SNPs." .format(input_Count_Col, input_Count_Row))
```
