## Supplemental Materials 1 for "Pronounced Genetic Separation Among Varieties of the *Primula cusickiana* Species Complex, a Great Basin Endemic": Austin_cartopy.html


In [2]:

```
import pandas as pd
```

In [36]:

```
# reading in Austin's file
k7 = pd.read_csv('K7_plot.csv', sep=' ')
print(k7.shape)
print(k7.columns)
print(k7.head())
```

```
(82, 10)
Index(['Group', 'Ind', 'PopID', 'P1', 'P2', 'P3', 'P4', 'P5', 'P6', 'P7'], dtype='object')
              Group  Ind PopID      P1      P2      P3      P4      P5  \
0  Cusickiana_Boise  Ck1   SRP  0.0352  0.0001  0.0000  0.0085  0.0008   
1  Cusickiana_Boise  Ck2   SRP  0.0378  0.0060  0.0002  0.0142  0.0013   
2  Cusickiana_Boise  Ck3   SRP  0.0347  0.0000  0.0000  0.0084  0.0000   
3  Cusickiana_Boise  Ck4   SRP  0.0378  0.0001  0.0000  0.0100  0.0013   
4  Cusickiana_Boise  Ck7   SRP  0.0396  0.0000  0.0000  0.0094  0.0002   

       P6      P7  
0  0.9548  0.0007  
1  0.9252  0.0154  
2  0.9560  0.0009  
3  0.9449  0.0059  
4  0.9506  0.0000
```

In [38]:

```
# need to make this file so that we can join more easliy with Austin's coord file
# spliting the "Group" column into to using the "_" as a split
k7[['Variety name', 'Site']]=k7['Group'].str.split('_',expand=True)
print(k7.shape)
print(k7.head())
```

```
(82, 12)
              Group  Ind PopID      P1      P2      P3      P4      P5  \
0  Cusickiana_Boise  Ck1   SRP  0.0352  0.0001  0.0000  0.0085  0.0008   
1  Cusickiana_Boise  Ck2   SRP  0.0378  0.0060  0.0002  0.0142  0.0013   
2  Cusickiana_Boise  Ck3   SRP  0.0347  0.0000  0.0000  0.0084  0.0000   
3  Cusickiana_Boise  Ck4   SRP  0.0378  0.0001  0.0000  0.0100  0.0013   
4  Cusickiana_Boise  Ck7   SRP  0.0396  0.0000  0.0000  0.0094  0.0002   

       P6      P7 Variety name   Site  
0  0.9548  0.0007   Cusickiana  Boise  
1  0.9252  0.0154   Cusickiana  Boise  
2  0.9560  0.0009   Cusickiana  Boise  
3  0.9449  0.0059   Cusickiana  Boise  
4  0.9506  0.0000   Cusickiana  Boise
```

In [41]:

```
# CR modified this file. Dropped cols not being used, and changed his "Site (Description)" to a more meaningful -
# I created the 'Site' column in which I selected the word that matches with what is now the 'Site' column in the k7 file above
#coord = pd.read_csv('Coordinates.csv', usecols = ['Variety name','PopName', 'latitude', 'longitude'])
coord = pd.read_csv('CR_Coordinates.csv')
print(coord.shape)
print(coord.columns)
print(coord.head())
```

```
(17, 5)
Index(['Variety name', 'PopName', 'Site', 'latitude', 'longitude'], dtype='object')
  Variety name PopName      Site   latitude   longitude
0   Cusickiana     SRP     Boise  43.684440 -116.187800
1   Cusickiana     SRP      Bear  44.981200 -116.680070
2   Cusickiana     SRP     Camas  43.310770 -115.091160
3   Cusickiana     SRP      CRMO  43.407805 -113.626270
4   Cusickiana     JAR  Jarbidge  41.958458 -115.315975
```

In [44]:

```
# merge the dataframes together
k7_coord = pd.merge(coord, k7, left_on=['Variety name', 'Site', 'PopName'],
                   right_on=['Variety name', 'Site', 'PopID'], how='right')

print(k7_coord.shape)
print(k7_coord.head())
```

```
(82, 15)
  Variety name PopName   Site  latitude  longitude             Group  Ind  \
0   Cusickiana     SRP  Boise  43.68444  -116.1878  Cusickiana_Boise  Ck1   
1   Cusickiana     SRP  Boise  43.68444  -116.1878  Cusickiana_Boise  Ck2   
2   Cusickiana     SRP  Boise  43.68444  -116.1878  Cusickiana_Boise  Ck3   
3   Cusickiana     SRP  Boise  43.68444  -116.1878  Cusickiana_Boise  Ck4   
4   Cusickiana     SRP  Boise  43.68444  -116.1878  Cusickiana_Boise  Ck7   

  PopID      P1      P2      P3      P4      P5      P6      P7  
0   SRP  0.0352  0.0001  0.0000  0.0085  0.0008  0.9548  0.0007  
1   SRP  0.0378  0.0060  0.0002  0.0142  0.0013  0.9252  0.0154  
2   SRP  0.0347  0.0000  0.0000  0.0084  0.0000  0.9560  0.0009  
3   SRP  0.0378  0.0001  0.0000  0.0100  0.0013  0.9449  0.0059  
4   SRP  0.0396  0.0000  0.0000  0.0094  0.0002  0.9506  0.0000
```

In [45]:

```
# saving this to a file
k7_coord.to_csv('CR_k7_coord.csv', index=False)
```

In [ ]:

```
# Second part
```

In [1]:

```
import pandas as pd
import matplotlib.patches as mpatches
import matplotlib.pyplot as plt
from scalebar import scale_bar # See section on scalebar.py
import cartopy.io.img_tiles as cimgt
import seaborn as sns
import cartopy.crs as ccrs
import cartopy.feature as cfeat
import matplotlib.pyplot as plt
from mpl_toolkits.axes_grid1.inset_locator import inset_axes
%matplotlib inline
```

In [69]:

```
k7_coord = pd.read_csv('CR_k7_coord.csv')
print(k7_coord.shape)
print(k7_coord.head())
```

```
(82, 15)
  Variety name PopName   Site  latitude  longitude             Group  Ind  \
0   Cusickiana     SRP  Boise  43.68444  -116.1878  Cusickiana_Boise  Ck1   
1   Cusickiana     SRP  Boise  43.68444  -116.1878  Cusickiana_Boise  Ck2   
2   Cusickiana     SRP  Boise  43.68444  -116.1878  Cusickiana_Boise  Ck3   
3   Cusickiana     SRP  Boise  43.68444  -116.1878  Cusickiana_Boise  Ck4   
4   Cusickiana     SRP  Boise  43.68444  -116.1878  Cusickiana_Boise  Ck7   

  PopID      P1      P2      P3      P4      P5      P6      P7  
0   SRP  0.0352  0.0001  0.0000  0.0085  0.0008  0.9548  0.0007  
1   SRP  0.0378  0.0060  0.0002  0.0142  0.0013  0.9252  0.0154  
2   SRP  0.0347  0.0000  0.0000  0.0084  0.0000  0.9560  0.0009  
3   SRP  0.0378  0.0001  0.0000  0.0100  0.0013  0.9449  0.0059  
4   SRP  0.0396  0.0000  0.0000  0.0094  0.0002  0.9506  0.0000
```

In [70]:

```
# parryi, cusickiana_Jarbidge, domensis, cusickiana_Owyhee, maguirei, cusickiana_SRP, nevadensis.


# Create a function to call the clusters with name that will corrspond with the pie chart in the map
def group_call(df):
    if (df['P7'] > 0.5):
        return "nevadensis"
    elif (df['P6'] > 0.5):
        return "cusickiana_SRP"
    elif (df['P5'] > 0.5):
        return "maguirei"    
    elif (df['P4'] > 0.5):
        return "cusickiana_Owyhee"
    elif (df['P3'] > 0.5):
        return "domensis"
    elif (df['P2'] > 0.5):
        return 'cusickiana_Jarbidge'
    elif (df['P1'] > 0.5):
        return "parryi"
    else:
        return "error"
```

In [71]:

```
# want the mean for each unique lat/long get the mean of each P# column
ll_grp = k7_coord.groupby(['latitude', 'longitude', 'PopID', 'Variety name'])[['P1', 'P2', 'P3', 'P4', 'P5', 'P6', 'P7']].mean()
ll_grp.reset_index(inplace=True)
# let's round some of the numbers to 4 digits
ll_grp2 = ll_grp.round(4)
print(ll_grp2)
print(ll_grp2.shape)
print(ll_grp2.columns)
print(ll_grp2)
```

```
    latitude  longitude PopID Variety name      P1      P2      P3      P4  \
0    38.3219  -115.4978   NEV   Nevadensis  0.0000  0.1921  0.1168  0.0199   
1    38.9123  -114.3087   NEV   Nevadensis  0.0003  0.0004  0.7668  0.1441   
2    39.1412  -113.3879   DOM     Domensis  0.0000  0.0000  0.9988  0.0003   
3    39.1429  -113.4056   DOM     Domensis  0.0003  0.0000  0.9983  0.0005   
4    40.5926  -115.3800   PAR       Parryi  0.9200  0.0000  0.0200  0.0201   
5    41.7427  -111.7617   MAG     Maguirei  0.0000  0.0002  0.0000  0.0007   
6    41.7454  -111.7523   MAG     Maguirei  0.0002  0.0001  0.0000  0.0000   
7    41.7771  -111.6328   MAG     Maguirei  0.0004  0.0000  0.0000  0.0004   
8    41.7983  -111.6424   MAG     Maguirei  0.0000  0.0000  0.0000  0.0000   
9    41.9585  -115.3160   JAR   Cusickiana  0.0000  0.9995  0.0000  0.0000   
10   42.1700  -117.6154   OWY   Cusickiana  0.0228  0.0868  0.0000  0.8693   
11   43.3108  -115.0912   SRP   Cusickiana  0.0005  0.0001  0.0000  0.0005   
12   43.4078  -113.6263   SRP   Cusickiana  0.0002  0.0002  0.0000  0.0002   
13   43.6844  -116.1878   SRP   Cusickiana  0.0370  0.0012  0.0000  0.0101   
14   44.9812  -116.6801   SRP   Cusickiana  0.0461  0.0017  0.0001  0.0119   

        P5      P6      P7  
0   0.0001  0.0008  0.6703  
1   0.0002  0.0009  0.0873  
2   0.0002  0.0002  0.0005  
3   0.0000  0.0001  0.0007  
4   0.0398  0.0000  0.0001  
5   0.9978  0.0000  0.0013  
6   0.9995  0.0000  0.0001  
7   0.9980  0.0002  0.0010  
8   1.0000  0.0000  0.0000  
9   0.0002  0.0002  0.0001  
10  0.0004  0.0001  0.0206  
11  0.0000  0.9966  0.0023  
12  0.0001  0.9985  0.0008  
13  0.0007  0.9463  0.0046  
14  0.0008  0.9346  0.0048  
(15, 11)
Index(['latitude', 'longitude', 'PopID', 'Variety name', 'P1', 'P2', 'P3',
       'P4', 'P5', 'P6', 'P7'],
      dtype='object')
    latitude  longitude PopID Variety name      P1      P2      P3      P4  \
0    38.3219  -115.4978   NEV   Nevadensis  0.0000  0.1921  0.1168  0.0199   
1    38.9123  -114.3087   NEV   Nevadensis  0.0003  0.0004  0.7668  0.1441   
2    39.1412  -113.3879   DOM     Domensis  0.0000  0.0000  0.9988  0.0003   
3    39.1429  -113.4056   DOM     Domensis  0.0003  0.0000  0.9983  0.0005   
4    40.5926  -115.3800   PAR       Parryi  0.9200  0.0000  0.0200  0.0201   
5    41.7427  -111.7617   MAG     Maguirei  0.0000  0.0002  0.0000  0.0007   
6    41.7454  -111.7523   MAG     Maguirei  0.0002  0.0001  0.0000  0.0000   
7    41.7771  -111.6328   MAG     Maguirei  0.0004  0.0000  0.0000  0.0004   
8    41.7983  -111.6424   MAG     Maguirei  0.0000  0.0000  0.0000  0.0000   
9    41.9585  -115.3160   JAR   Cusickiana  0.0000  0.9995  0.0000  0.0000   
10   42.1700  -117.6154   OWY   Cusickiana  0.0228  0.0868  0.0000  0.8693   
11   43.3108  -115.0912   SRP   Cusickiana  0.0005  0.0001  0.0000  0.0005   
12   43.4078  -113.6263   SRP   Cusickiana  0.0002  0.0002  0.0000  0.0002   
13   43.6844  -116.1878   SRP   Cusickiana  0.0370  0.0012  0.0000  0.0101   
14   44.9812  -116.6801   SRP   Cusickiana  0.0461  0.0017  0.0001  0.0119   

        P5      P6      P7  
0   0.0001  0.0008  0.6703  
1   0.0002  0.0009  0.0873  
2   0.0002  0.0002  0.0005  
3   0.0000  0.0001  0.0007  
4   0.0398  0.0000  0.0001  
5   0.9978  0.0000  0.0013  
6   0.9995  0.0000  0.0001  
7   0.9980  0.0002  0.0010  
8   1.0000  0.0000  0.0000  
9   0.0002  0.0002  0.0001  
10  0.0004  0.0001  0.0206  
11  0.0000  0.9966  0.0023  
12  0.0001  0.9985  0.0008  
13  0.0007  0.9463  0.0046  
14  0.0008  0.9346  0.0048
```

In [82]:

```
# now apply the group_call function
ll_grp2['Group'] = ll_grp.apply(group_call, axis=1)
print(ll_grp2.head())
```

```
   latitude  longitude PopID Variety name      P1      P2      P3      P4  \
0   38.3219  -115.4978   NEV   Nevadensis  0.0000  0.1921  0.1168  0.0199   
1   38.9123  -114.3087   NEV   Nevadensis  0.0003  0.0004  0.7668  0.1441   
2   39.1412  -113.3879   DOM     Domensis  0.0000  0.0000  0.9988  0.0003   
3   39.1429  -113.4056   DOM     Domensis  0.0003  0.0000  0.9983  0.0005   
4   40.5926  -115.3800   PAR       Parryi  0.9200  0.0000  0.0200  0.0201   

       P5      P6      P7       Group  
0  0.0001  0.0008  0.6703  nevadensis  
1  0.0002  0.0009  0.0873    domensis  
2  0.0002  0.0002  0.0005    domensis  
3  0.0000  0.0001  0.0007    domensis  
4  0.0398  0.0000  0.0001      parryi
```

In [95]:

```
ll_grp2['Group'].value_counts()
```

Out[95]:

```
maguirei               4
cusickiana_SRP         4
domensis               3
parryi                 1
cusickiana_Jarbidge    1
nevadensis             1
cusickiana_Owyhee      1
Name: Group, dtype: int64
```

In [93]:

```
# save this too
ll_grp2.to_csv('CR_k7_mean_with_latlon.csv', index=False)
```

In [112]:

```
# transition back again
ll_grp2 = pd.read_csv('CR_k7_mean_with_latlon.csv')
print(ll_grp2.shape)
```

```
(15, 12)
```

In [113]:

```
# Austin wants two pies removed b/c one is pretty much located on top of the other and also look the same
# remove row 5, maguirei (Greenhouse wall)
# remove row 2, domensis (Sawtooth Canyon)
ll_grp2.drop([2,5],0,inplace=True)
# So there is no chance of problems down-stream when making the map I'll reset the index
ll_grp2.reset_index(inplace=True, drop=True)

print(ll_grp2)
```

```
    latitude  longitude PopID Variety name      P1      P2      P3      P4  \
0    38.3219  -115.4978   NEV   Nevadensis  0.0000  0.1921  0.1168  0.0199   
1    38.9123  -114.3087   NEV   Nevadensis  0.0003  0.0004  0.7668  0.1441   
2    39.1429  -113.4056   DOM     Domensis  0.0003  0.0000  0.9983  0.0005   
3    40.5926  -115.3800   PAR       Parryi  0.9200  0.0000  0.0200  0.0201   
4    41.7454  -111.7523   MAG     Maguirei  0.0002  0.0001  0.0000  0.0000   
5    41.7771  -111.6328   MAG     Maguirei  0.0004  0.0000  0.0000  0.0004   
6    41.7983  -111.6424   MAG     Maguirei  0.0000  0.0000  0.0000  0.0000   
7    41.9585  -115.3160   JAR   Cusickiana  0.0000  0.9995  0.0000  0.0000   
8    42.1700  -117.6154   OWY   Cusickiana  0.0228  0.0868  0.0000  0.8693   
9    43.3108  -115.0912   SRP   Cusickiana  0.0005  0.0001  0.0000  0.0005   
10   43.4078  -113.6263   SRP   Cusickiana  0.0002  0.0002  0.0000  0.0002   
11   43.6844  -116.1878   SRP   Cusickiana  0.0370  0.0012  0.0000  0.0101   
12   44.9812  -116.6801   SRP   Cusickiana  0.0461  0.0017  0.0001  0.0119   

        P5      P6      P7                Group  
0   0.0001  0.0008  0.6703           nevadensis  
1   0.0002  0.0009  0.0873             domensis  
2   0.0000  0.0001  0.0007             domensis  
3   0.0398  0.0000  0.0001               parryi  
4   0.9995  0.0000  0.0001             maguirei  
5   0.9980  0.0002  0.0010             maguirei  
6   1.0000  0.0000  0.0000             maguirei  
7   0.0002  0.0002  0.0001  cusickiana_Jarbidge  
8   0.0004  0.0001  0.0206    cusickiana_Owyhee  
9   0.0000  0.9966  0.0023       cusickiana_SRP  
10  0.0001  0.9985  0.0008       cusickiana_SRP  
11  0.0007  0.9463  0.0046       cusickiana_SRP  
12  0.0008  0.9346  0.0048       cusickiana_SRP
```

In [114]:

```
# just checking all looks correct - should now be 13 rows
print(ll_grp2.shape)
```

```
(13, 12)
```

In [74]:

```
# need 7 colors for the 7 legend names
colors = ["#2171B5","#D95F02","#7570B3","#E7298A","#66A61E","#8C510A","#666666"]
len(colors) # 7 colors
```

Out[74]:

```
7
```

In [92]:

```
# legend names
P_names = ['parryi', 'cusickiana_Jarbidge', 'domensis', 'cusickiana_Owyhee', 'maguirei', 'cusickiana_SRP', 'nevadensis']
```

In [77]:

```
# Settting up for the legend
# create a patch object for associating name with corresponding color for the legend
my_patches = []
for i in range(len(P_names)):
    this_patch = mpatches.Patch(color=colors[i], label=P_names[i])
    my_patches.append(this_patch)

print(len(my_patches)) # 7
```

```
7
```

In [89]:

```
# Get map boundaries. 
# Note that I add/sub from lat/long so map boundary goesbeyond the plotted points.
# This is so the point locations are not on the edge of the map
west_long = float( (ll_grp2['longitude'].min()) + 7 )
east_long = float( (ll_grp2['longitude'].max()) - 7 )
north_lat = float( (ll_grp2['latitude'].max()) + 0.8 )
south_lat = float( (ll_grp2['latitude'].min()) - 0.8 )

print(west_long)
print(east_long)
print(north_lat)
print(south_lat)
```

```
-110.6154
-118.6328
45.7812
37.5219
```

In [115]:

```
# creating these lists for the map we will draw
LAT = ll_grp2['latitude'].values.tolist()
LONG = ll_grp2['longitude'].values.tolist()
R1 = ll_grp2['P1'].values.tolist()
R2 = ll_grp2['P2'].values.tolist()
R3 = ll_grp2['P3'].values.tolist()
R4 = ll_grp2['P4'].values.tolist()
R5 = ll_grp2['P5'].values.tolist()
R6 = ll_grp2['P6'].values.tolist()
R7 = ll_grp2['P7'].values.tolist()
```

In [116]:

```
# making a list of lists for lat/long and for the P# columns
# the lat/long combo (LatLong_list) is for where we will plot the pie graph
# the P# columns combo (Wedge_list) is for the size of each pie wedge

# Create an empty list 
LatLong_list =[]
Wedge_list = []
  
# Iterate over each row 
for index, rows in ll_grp2.iterrows(): 
    # Create list for the current row 
    my_latlong =[rows['longitude'], rows['latitude'] ]
    my_wedges = [rows['P1'], rows['P2'], rows['P3'], rows['P4'], rows['P5'], rows['P6'], rows['P7']]
      
    # append the list to the final list 
    LatLong_list.append(my_latlong)
    Wedge_list.append(my_wedges)

# Print the list just so you can see
print(Row_list)
print(Wedge_list)
```

```
[[-115.4978, 38.3219], [-114.3087, 38.9123], [-113.3879, 39.1412], [-113.4056, 39.1429], [-115.38, 40.5926], [-111.7617, 41.7427], [-111.7523, 41.7454], [-111.6328, 41.7771], [-111.6424, 41.7983], [-115.316, 41.9585], [-117.6154, 42.17], [-115.0912, 43.3108], [-113.6263, 43.4078], [-116.1878, 43.6844], [-116.6801, 44.9812]]
[[0.0, 0.1921, 0.1168, 0.0199, 0.0001, 0.0008, 0.6703], [0.0003, 0.0004, 0.7668, 0.1441, 0.0002, 0.0009, 0.0873], [0.0003, 0.0, 0.9983, 0.0005, 0.0, 0.0001, 0.0007], [0.92, 0.0, 0.02, 0.0201, 0.0398, 0.0, 0.0001], [0.0002, 0.0001, 0.0, 0.0, 0.9995, 0.0, 0.0001], [0.0004, 0.0, 0.0, 0.0004, 0.998, 0.0002, 0.001], [0.0, 0.0, 0.0, 0.0, 1.0, 0.0, 0.0], [0.0, 0.9995, 0.0, 0.0, 0.0002, 0.0002, 0.0001], [0.0228, 0.0868, 0.0, 0.8693, 0.0004, 0.0001, 0.0206], [0.0005, 0.0001, 0.0, 0.0005, 0.0, 0.9966, 0.0023], [0.0002, 0.0002, 0.0, 0.0002, 0.0001, 0.9985, 0.0008], [0.037000000000000005, 0.0012, 0.0, 0.0101, 0.0007, 0.9463, 0.0046], [0.0461, 0.0017, 0.0001, 0.0119, 0.0008, 0.9346, 0.0048]]
```

In [101]:

```
# reminder of the dataframe
print(ll_grp2)
```

```
    index  latitude  longitude PopID Variety name      P1      P2      P3  \
0       0   38.3219  -115.4978   NEV   Nevadensis  0.0000  0.1921  0.1168   
1       1   38.9123  -114.3087   NEV   Nevadensis  0.0003  0.0004  0.7668   
2       3   39.1429  -113.4056   DOM     Domensis  0.0003  0.0000  0.9983   
3       4   40.5926  -115.3800   PAR       Parryi  0.9200  0.0000  0.0200   
4       6   41.7454  -111.7523   MAG     Maguirei  0.0002  0.0001  0.0000   
5       7   41.7771  -111.6328   MAG     Maguirei  0.0004  0.0000  0.0000   
6       8   41.7983  -111.6424   MAG     Maguirei  0.0000  0.0000  0.0000   
7       9   41.9585  -115.3160   JAR   Cusickiana  0.0000  0.9995  0.0000   
8      10   42.1700  -117.6154   OWY   Cusickiana  0.0228  0.0868  0.0000   
9      11   43.3108  -115.0912   SRP   Cusickiana  0.0005  0.0001  0.0000   
10     12   43.4078  -113.6263   SRP   Cusickiana  0.0002  0.0002  0.0000   
11     13   43.6844  -116.1878   SRP   Cusickiana  0.0370  0.0012  0.0000   
12     14   44.9812  -116.6801   SRP   Cusickiana  0.0461  0.0017  0.0001   

        P4      P5      P6      P7                Group  
0   0.0199  0.0001  0.0008  0.6703           nevadensis  
1   0.1441  0.0002  0.0009  0.0873             domensis  
2   0.0005  0.0000  0.0001  0.0007             domensis  
3   0.0201  0.0398  0.0000  0.0001               parryi  
4   0.0000  0.9995  0.0000  0.0001             maguirei  
5   0.0004  0.9980  0.0002  0.0010             maguirei  
6   0.0000  1.0000  0.0000  0.0000             maguirei  
7   0.0000  0.0002  0.0002  0.0001  cusickiana_Jarbidge  
8   0.8693  0.0004  0.0001  0.0206    cusickiana_Owyhee  
9   0.0005  0.0000  0.9966  0.0023       cusickiana_SRP  
10  0.0002  0.0001  0.9985  0.0008       cusickiana_SRP  
11  0.0101  0.0007  0.9463  0.0046       cusickiana_SRP  
12  0.0119  0.0008  0.9346  0.0048       cusickiana_SRP
```

In [123]:

```
# We have noticed that the three green pies for maguirei overlap each other.
# Want to offset these pies so we can see them individually
# creating off_set lists for position 4,5,6

#LAT_offset = [0,0,0, 41.7454,41.7771,41.7983, 0,0,0,0,0,0,0]
#LONG_offset = [0,0,0, -111.7523,-111.6328,-111.6424, 0,0,0,0,0,0,0]


#LAT_offset = [0,0,0,0, 41.7454,41.7771,41.96, 0,0,0,0,0,0]
#LONG_offset = [0,0,0,0, -111.95,-111.33,-111.6424, 0,0,0,0,0,0]
LAT_offset = [0,0,0,0, 41.7454,41.7771,42.0, 0,0,0,0,0,0]
LONG_offset = [0,0,0,0, -112.0,-111.33,-111.6424, 0,0,0,0,0,0]
```

In [124]:

```
# Make the map
# First pull in a greyscale topography map from World_Terrain_Base website
arcgis_url = 'https://server.arcgisonline.com/ArcGIS/rest/services/World_Terrain_Base/MapServer/tile/{z}/{y}/{x}.jpg'
tiles = cimgt.GoogleTiles(url=arcgis_url)

# Now set up figure and axis for map
fig = plt.figure(figsize=(12,12))
ax = fig.add_subplot(1,1,1, projection=ccrs.LambertConformal())

# Set the lat/long dimensions for the map
ax.set_extent([west_long, east_long, south_lat, north_lat], crs = ccrs.PlateCarree())

# Add features
ax.add_feature(cfeat.OCEAN.with_scale('50m'), alpha=0.8, zorder=2)
ax.add_feature(cfeat.BORDERS.with_scale('50m'), linewidth=0.4, alpha=0.8)
ax.add_feature(cfeat.STATES.with_scale('10m'), linewidth=1.0, alpha=0.3)
ax.add_feature(cfeat.RIVERS.with_scale('10m'), linewidth=0.8, alpha=0.8, zorder=5)
ax.add_feature(cfeat.LAKES.with_scale('10m'), alpha=0.8, zorder=4)

# Now add the topomap from the arcgis_url above
ax.add_image(tiles,8, cmap='gray')

# Add the scale bar.
# Use the numbers in parentheses to move scale bar on x,y axis
# The 5_00 designates scale bar to be at 500 km. You can change this.
scale_bar(ax, (0.6, 0.05), 1_00, color='black', zorder=6)

# here, you can change size of pie chart with the width value entered when calling the function
def plot_pie_inset(data,ilon,ilat,ax,width):
    #inset_axes allows us to plot over the map
    ax_sub= inset_axes(ax, width=width, height=width, loc=10, 
                       bbox_to_anchor=(ilon, ilat),
                       bbox_transform=ax.transData, 
                       borderpad=0)
    # here, drawing the pie with associated colors
    # see matplotlib.pyplot.pie for more info
    wedges,texts= ax_sub.pie(data, colors=colors, wedgeprops={'edgecolor' :'black', 'linewidth': 0.6})
    # making sure all adds up to 100% so get full circle for the pie.
    ax_sub.set_aspect("equal")

    
for row in range(ll_grp2.shape[0]):
    # for each row in the dataframe, transform the lat/long
    lonr,latr =  ccrs.LambertConformal().transform_point(LONG[row],LAT[row], ccrs.PlateCarree())
    if LAT_offset[row] > 0:
        lonr_offset,latr_offset =  ccrs.LambertConformal().transform_point(LONG_offset[row],LAT_offset[row], ccrs.PlateCarree())
        ax.plot([lonr, lonr_offset], [latr, latr_offset], color='k', lw=1, zorder=2,marker='o', markersize=0.3)
        plot_pie_inset(Wedge_list[row],lonr_offset,latr_offset,ax,0.3)
    else:
        # now draw the pie. Wedge_list[row] is a list of the proportions for the pie wedges
        plot_pie_inset(Wedge_list[row],lonr,latr,ax,0.3)

# Add the legend
ax.legend(handles=my_patches, loc='upper left')

# Save the plot. Can change the dpi for higher/lower resolution
plt.savefig("CR_Austin_relief_map.pdf",bbox_inches='tight', dpi=700)
```

In [ ]:
