## Supplemental Materials 1 for "Pronounced Genetic Separation Among Varieties of the *Primula cusickiana* Species Complex, a Great Basin Endemic": scalebar.py

```
import numpy as np
import cartopy.crs as ccrs
import cartopy.geodesic as cgeo

# Much thanks to whowver wrote this python script. I got it off the internet:
# https://stackoverflow.com/questions/32333870/how-can-i-show-a-km-ruler-on-a-cartopy-matplotlib-plot

def _axes_to_lonlat(ax, coords):
    """(lon, lat) from axes coordinates."""
    display = ax.transAxes.transform(coords)
    data = ax.transData.inverted().transform(display)
    lonlat = ccrs.PlateCarree().transform_point(*data, ax.projection)

    return lonlat


def _upper_bound(start, direction, distance, dist_func):
    """A point farther than distance from start, in the given direction.

    It doesn't matter which coordinate system start is given in, as long
    as dist_func takes points in that coordinate system.

    Args:
        start:     Starting point for the line.
        direction  Nonzero (2, 1)-shaped array, a direction vector.
        distance:  Positive distance to go past.
        dist_func: A two-argument function which returns distance.

    Returns:
        Coordinates of a point (a (2, 1)-shaped NumPy array).
    """
    if distance <= 0:
        raise ValueError(f"Minimum distance is not positive: {distance}")

    if np.linalg.norm(direction) == 0:
        raise ValueError("Direction vector must not be zero.")

    # Exponential search until the distance between start and end is
    # greater than the given limit.
    length = 0.1
    end = start + length * direction

    while dist_func(start, end) < distance:
        length *= 2
        end = start + length * direction

    return end


def _distance_along_line(start, end, distance, dist_func, tol):
    """Point at a distance from start on the segment  from start to end.

    It doesn't matter which coordinate system start is given in, as long
    as dist_func takes points in that coordinate system.

    Args:
        start:     Starting point for the line.
        end:       Outer bound on point's location.
        distance:  Positive distance to travel.
        dist_func: Two-argument function which returns distance.
        tol:       Relative error in distance to allow.

    Returns:
        Coordinates of a point (a (2, 1)-shaped NumPy array).
    """
    initial_distance = dist_func(start, end)
    if initial_distance < distance:
        raise ValueError(f"End is closer to start ({initial_distance}) than "
                         f"given distance ({distance}).")

    if tol <= 0:
        raise ValueError(f"Tolerance is not positive: {tol}")

    # Binary search for a point at the given distance.
    left = start
    right = end

    while not np.isclose(dist_func(start, right), distance, rtol=tol):
        midpoint = (left + right) / 2

        # If midpoint is too close, search in second half.
        if dist_func(start, midpoint) < distance:
            left = midpoint
        # Otherwise the midpoint is too far, so search in first half.
        else:
            right = midpoint

    return right


def _point_along_line(ax, start, distance, angle=0, tol=0.01):
    """Point at a given distance from start at a given angle.

    Args:
        ax:       CartoPy axes.
        start:    Starting point for the line in axes coordinates.
        distance: Positive physical distance to travel.
        angle:    Anti-clockwise angle for the bar, in radians. Default: 0
        tol:      Relative error in distance to allow. Default: 0.01

    Returns:
        Coordinates of a point (a (2, 1)-shaped NumPy array).
    """
    # Direction vector of the line in axes coordinates.
    direction = np.array([np.cos(angle), np.sin(angle)])

    geodesic = cgeo.Geodesic()

    # Physical distance between points.
    def dist_func(a_axes, b_axes):
        a_phys = _axes_to_lonlat(ax, a_axes)
        b_phys = _axes_to_lonlat(ax, b_axes)

        # Geodesic().inverse returns a NumPy MemoryView like [[distance,
        # start azimuth, end azimuth]].
        return geodesic.inverse(a_phys, b_phys).base[0, 0]

    end = _upper_bound(start, direction, distance, dist_func)

    return _distance_along_line(start, end, distance, dist_func, tol)


def scale_bar(ax, location, length, metres_per_unit=1000, unit_name='km',
              tol=0.01, angle=0, color='black', linewidth=3, text_offset=0.005,
              ha='center', va='bottom', plot_kwargs=None, text_kwargs=None,
              **kwargs):
    """Add a scale bar to CartoPy axes.

    For angles between 0 and 90 the text and line may be plotted at
    slightly different angles for unknown reasons. To work around this,
    override the 'rotation' keyword argument with text_kwargs.

    Args:
        ax:              CartoPy axes.
        location:        Position of left-side of bar in axes coordinates.
        length:          Geodesic length of the scale bar.
        metres_per_unit: Number of metres in the given unit. Default: 1000
        unit_name:       Name of the given unit. Default: 'km'
        tol:             Allowed relative error in length of bar. Default: 0.01
        angle:           Anti-clockwise rotation of the bar.
        color:           Color of the bar and text. Default: 'black'
        linewidth:       Same argument as for plot.
        text_offset:     Perpendicular offset for text in axes coordinates.
                         Default: 0.005
        ha:              Horizontal alignment. Default: 'center'
        va:              Vertical alignment. Default: 'bottom'
        **plot_kwargs:   Keyword arguments for plot, overridden by **kwargs.
        **text_kwargs:   Keyword arguments for text, overridden by **kwargs.
        **kwargs:        Keyword arguments for both plot and text.
    """
    # Setup kwargs, update plot_kwargs and text_kwargs.
    if plot_kwargs is None:
        plot_kwargs = {}
    if text_kwargs is None:
        text_kwargs = {}

    plot_kwargs = {'linewidth': linewidth, 'color': color, **plot_kwargs,
                   **kwargs}
    text_kwargs = {'ha': ha, 'va': va, 'rotation': angle, 'color': color,
                   **text_kwargs, **kwargs}

    # Convert all units and types.
    location = np.asarray(location)  # For vector addition.
    length_metres = length * metres_per_unit
    angle_rad = angle * np.pi / 180

    # End-point of bar.
    end = _point_along_line(ax, location, length_metres, angle=angle_rad,
                            tol=tol)

    # Coordinates are currently in axes coordinates, so use transAxes to
    # put into data coordinates. *zip(a, b) produces a list of x-coords,
    # then a list of y-coords.
    ax.plot(*zip(location, end), transform=ax.transAxes, **plot_kwargs)

    # Push text away from bar in the perpendicular direction.
    midpoint = (location + end) / 2
    offset = text_offset * np.array([-np.sin(angle_rad), np.cos(angle_rad)])
    text_location = midpoint + offset

    # 'rotation' keyword argument is in text_kwargs.
    ax.text(*text_location, f"{length} {unit_name}", rotation_mode='anchor',
            transform=ax.transAxes, **text_kwargs)
```
