## Supplemental Materials 1 for "Pronounced Genetic Separation Among Varieties of the *Primula cusickiana* Species Complex, a Great Basin Endemic": DAPC_code.R

```
library(adegenet)
# %%% DAPC--JUNESUBSET STRUCTURE FILE %%% ----
setwd("/home/akoontz11/kaiser/Code/ipyrad/JuneSubset/Outfiles")

# Read in STRUCTURE file to genind object
# 82 individuals, 1277 loci, with genotype labels. Genotypes NOT coded by a single row
junesub.genind <- import2genind("JuneSubset.str")
str(junesub.genind)

# Generate color palette 
colors <- c("#2171B5","#D95F02","#7570B3","#E7298A","#66A61E","#8C510A","#666666")
show_col(colors)

# Find number of clusters
grp <- find.clusters(junesub.genind, max.n.clust=60)
# Number of retained PCs (here, there is no cost to retaining a lot of PCs): 80
# Number of clusters: 7 (even though 11 has the lowest BIC)
grp$stat

# Conduct DAPC
dapc1 <- dapc(x=junesub.genind, pop=grp$grp)
# Number of retained PCs: 7
# Number of discriminant functions to retain: 2
str(dapc1)
# Export DAPC object
saveRDS(dapc1, "DAPC.RDS")
# List grouping of each individual (helpful for determining correct coloration)
dapc1$grp

# Make a scatterplot of initial DAPC
scatter(dapc1)
scatter(dapc1, label = NULL)
scatter(dapc1, clab = 0.5, label = letters[1:15])
scatter(dapc1, clab = 0.5, scree.da = F)

# Original DAPC image
scatter(dapc1, scree.da=F, bg="white", pch=20, cell=0, cstar=0, solid=0.6, clab=0, legend=T,
        posi.leg=locator(n=1), cleg=1.0, cex=1.8, inset.solid=1, 
        col = c("#66A61E","#E7298A","#7570B3","#2171B5","#666666","#8C510A","#D95F02"), 
        txt.leg = c("maguirei","cusickiana_Owyhee","domensis","parryi","nevadensis","cusickiana_Idaho","cusickiana_Jarbidge"))

# Systematic Botany figure
txt.leg <- c("cusickiana_Idaho","domensis","parryi","cusickiana_Owyhee","cusickiana_Jarbidge","nevadensis","maguirei")
par(mar=c(5,1,0,1)+0.1, mgp = c(3,1,1))
scatter(dapc1, clab = 0, scree.da = F, cstar=0, solid=0.6, cex=1.2,
        col=c("#8C510A","#7570B3","#2171B5","#E7298A","#D95F02","#666666","#66A61E"))
legend(x=-55, y=1.0, legend=txt.leg, cex=1, ncol=2, pch=16, bty="n",
       col=c("#8C510A","#7570B3","#2171B5","#E7298A","#D95F02","#666666","#66A61E"),
       x.intersp=0.2, y.intersp=0.3, text.width=20, xjust=0)
text(x=-63, y=1, cex=0.8, "PC 1: 49.04%")
text(x=-1, y=22, cex=0.8, srt=90, "PC 2: 27.27%")

# Plot for ESA
scatter(dapc1, scree.da=F, bg="white", pch=20, cell=0, cstar=0, solid=0.6, clab=0, legend=F,
        cleg=1.0, cex=2.5, inset.solid=1, 
        col = c("#E7298A","#D95F02","#8C510A","#66A61E","#666666","#2171B5","#7570B3"))

# %%% DAPC--COMPLEX ONLY %%% ----
setwd("/home/akoontz11/kaiser/Code/ipyrad/JuneSubset/Outfiles")

# Read in STRUCTURE file to genind object
# 75 individuals, 1277 loci, with genotype labels. Genotypes NOT coded by a single row
complexVars.genind <- import2genind("JuneSubset_ComplexOnly.str")
str(complexVars.genind)

# Generate color palette 
colors7 <- c("#2171B5","#D95F02","#7570B3","#E7298A","#66A61E","#8C510A","#666666")
colors6 <- c("#D95F02","#7570B3","#E7298A","#66A61E","#8C510A","#666666")
show_col(colors7)

# 6 Clusters
# Find number of clusters
grp2 <- find.clusters(complexVars.genind, max.n.clust=60)
# Number of retained PCs (here, there is no cost to retaining a lot of PCs): 80
# Number of clusters: 6 
grp2$stat

# Conduct DAPC
dapc6 <- dapc(x=complexVars.genind, pop=grp2$grp)
# Number of retained PCs: 6
# Number of discriminant functions to retain: 4
str(dapc6)
# List grouping of each individual (helpful for determining correct coloration)
dapc6$grp
# PC axis variance values
eig.perc <- 100*dapc6$eig/sum(dapc6$eig)

# Plotting
txt.leg <- c("maguirei","cusickiana_SRP","domensis","cusickiana_Owyhee","nevadensis_GRBA","cusickiana_Jarbidge")
par(mar=c(5,1,0,1)+0.1, mgp = c(3,1,1))
scatter(dapc6, clab = 0, scree.da = F, cstar=0, solid=0.6, cex=1.2, 
        col=c("#66A61E","#8C510A","#7570B3","#E7298A","#666666","#D95F02"))
legend(x=1, y=35, legend=txt.leg, cex=0.8, ncol=2, pch=16, bty="n",
       col=c("#66A61E","#8C510A","#7570B3","#E7298A","#666666","#D95F02"),
       x.intersp=0.25, y.intersp=0.3, text.width=30, xjust=0)
text(x=40, y=1, cex=0.8, "PC 1: 50.98%")
text(x=-1, y=13, cex=0.8, srt=90, "PC 2: 23.29%")

# 7 Clusters
# Find number of clusters
grp2 <- find.clusters(complexVars.genind, max.n.clust=60)
# Number of retained PCs (here, there is no cost to retaining a lot of PCs): 80
# Number of clusters: 7 
grp2$stat

# Conduct DAPC
dapc7 <- dapc(x=complexVars.genind, pop=grp2$grp)
# Number of retained PCs: 7
# Number of discriminant functions to retain: 5
str(dapc7)
# List grouping of each individual (helpful for determining correct coloration)
dapc7$grp
# PC axis variance values
eig.perc <- 100*dapc7$eig/sum(dapc7$eig)

# Plotting
txt.leg <- c("cusickiana_Jarbidge","nevadensis_Troy","maguirei","cusickiana_IdahoSouth","cusickiana_Owyhee","cusickiana_IdahoNorth","domensis_nevadensisGRBA")
par(mar=c(5,1,0,1)+0.1, mgp = c(3,1,1))
scatter(dapc7, clab = 0, scree.da = F, cstar=0, solid=0.6, cex=1.2, 
        col=c("#D95F02","#666666","#66A61E","#2171B5","#E7298A","#8C510A","#7570B3"))
legend(x=-70, y=2.0, legend=txt.leg, cex=1, ncol=2, pch=16, bty="n",
       col=c("#D95F02","#666666","#66A61E","#2171B5","#E7298A","#8C510A","#7570B3"),
       x.intersp=0.2, y.intersp=0.3, text.width=20, xjust=0)
text(x=40, y=1, cex=0.8, "PC 1: 45.18%")
text(x=-1, y=15, cex=0.8, srt=90, "PC 2: 27.52%")
```
