## Supplemental Materials 1 for "Pronounced Genetic Separation Among Varieties of the *Primula cusickiana* Species Complex, a Great Basin Endemic": JaccardDistances.R

```
# Code for plotting a histogram of Jaccard distances, from vcf2Jaccard.py output

# Initial clustering diagram----
# Read in and reformat the data
setwd("/home/austin/Documents/WolfLab/Primula/Analysis/ipyrad/ipyrad_UniqueReps/UniqueReps_outfiles/vcf2Jaccard_output/")
JMat <- read.table("Jaccob_sim_means.csv", header=TRUE, sep = ",")
# Transpose rows and columns to get Jaccard similarities for each variety in a column
JMat <- t(as.matrix(JMat))
# Get rid of 1st row that just contains variety names
JMat <- JMat[-1,]
# Capture sample names
sample.names <- rownames(JMat)
# Change strings of numbers into numeric
JMat <- apply(JMat,2,as.numeric)
# Assign row/column names
rownames(JMat) <- sample.names ; colnames(JMat) <- sample.names

# Find matrix values corresponding to replicates matching their "originals"
rep_positions <- grep("_R",colnames(JMat))
comps_positions <- rep_positions -1
indices <- cbind(rep_positions,comps_positions)
indices
JMat[indices]

which(JMat >= 0.98,arr.ind=TRUE)
JMat[90:96,90:96]

# Basic histogram...
hist(JMat, xlab="Jaccard distance value", ylab="Number of pairwise comparisons", main="Min Samples per Locus: 10",
     breaks=seq(from=0, to=1, by=0.02),xlim=c(0,1.0))
abline(v=JMat[indices],col=550)

hist(JMat, xlab="Jaccard distance value", ylab="Number of pairwise comparisons", main="Complex-Wide Jaccard Similarities",
     breaks=seq(from=0, to=1, by=0.02),xlim=c(0,1.0), mgp=c(3,1,0), mar=c(5,4,1,1)+0.1, yaxt="n")
axis(side=2, at=seq(0,800,by=200), labels=T, tick=T, line=-1, ylab="Number of pairwise comparisons")
abline(v=JMat[indices],col="red")

# Zoomed in
hist(JMat, xlab="Jaccard distance value", ylab="Number of pairwise comparisons", main="Jaccard distances: replicates",
     breaks=seq(from=0, to=1, by=0.01),xlim=c(0.9,1.0))
abline(v=JMat[indices],col=550)
text(0.9877529, 500, label="321")

# Hierarchical clustering tree
mat <- 1 - JMat
mat[upper.tri(mat)] <- t(mat[lower.tri(mat)])
mat[is.na(mat)] <- median(mat, na.rm=TRUE)

plot(hclust(as.dist(mat)), main="Primula cusickiana species complex")

# PCA, using eigen
spec.decomp <- eigen(JMat) # Error: infinite or mising values in x
spec.decomp <- eigen(mat) # Inverse of Jmat works...
plot(spec.decomp$vectors)
plot(spec.decomp$values)
```
