## Supplemental Materials 1 for "Pronounced Genetic Separation Among Varieties of the *Primula cusickiana* Species Complex, a Great Basin Endemic": LociVSIndividuals.R

```
# Code for plotting individuals for a given number of loci versus number of loci

loci.n.data <- read.table("Loci_per_Individual.txt",header=TRUE)
loci.n.data

indv.per.locus <- loci.n.data$samples/loci.n.data$locus_coverage
indv.per.locus[which(is.infinite(indv.per.locus))] <- 0

plot(indv.per.locus ~ loci.n.data$locus_coverage, ylab = "Individuals per locus", xlab = "Number of loci retrieved")
```
