## Supplemental Materials 1 for "Pronounced Genetic Separation Among Varieties of the *Primula cusickiana* Species Complex, a Great Basin Endemic": ReadCounts.R

```
# Script comparing the number of raw reads to filtered reads, for the UniqueReps ipyrad run.

library(plotrix)

# Read in and reformat the data (derived from ipyrad output text file)
setwd("/home/akoontz11/kaiser/Code/ipyrad/UniqueReps/Outfiles_UniqueReps/")
data <- read.table("UniqueReps_Reads.txt", header=TRUE)
data$SampleName <- as.character(data$SampleName)
str(data)

# ---- RAW READS ----
# Collect information on raw reads
raw_reads_mean <- round(mean(data$reads_raw),digits = 2)
raw_reads_sd <- round(sd(data$reads_raw),digits = 2)
raw_reads_se <- round(sd(data$reads_raw)/sqrt(sum(!is.na(data$reads_raw))),digits = 2)
std.error(data$reads_raw)

# Plot raw reads information for each sample
barplot(height=data$reads_raw, xlab="SampleID", ylab="Number of reads", main="Raw reads per sample", xaxs="i",
        names.arg=data$SampleNames, las=2, space=0.7, width=3.2, xlim=c(0,475), cex.names=0.7, mgp=c(3.2,0.4,0))
mtext(paste0("Mean = ",raw_reads_mean,"   Standard error = ",raw_reads_se))

# Plot a histogram of raw reads
hist(x=data$reads_raw, breaks=seq(from=0, to=7500000, by=500000), xlab="Number of reads per sample", ylab="Number of samples", main="Raw read histogram",
     xlim=c(0,6000000), lab=c(10,9,7))

# ---- FILTERED READS ----
# Collect information on filtered reads
filtered_reads_mean <- round(mean(data$reads_passed_filter),digits = 2)
filtered_reads_sd <- round(sd(data$reads_passed_filter),digits = 2)
filtered_reads_se <- round(sd(data$reads_passed_filter)/sqrt(sum(!is.na(data$reads_passed_filter))),digits = 2)
std.error(data$reads_passed_filter)

# Plot filtered reads information for each sample
barplot(height=data$reads_passed_filter, xlab="SampleID", ylab="Number of reads", main="Filtered reads per sample", xaxs="i",
        names.arg=data$SampleNames, las=2, space=0.7, width=3.2, xlim=c(0,475), cex.names=0.7, mgp=c(3.2,0.4,0))
mtext(paste0("Mean = ",filtered_reads_mean,"   Standard error = ",filtered_reads_se))

# Plot a histogram of filtered reads
hist(x=data$reads_passed_filter, breaks=seq(from=0, to=4000000, by=250000), xlab="Number of reads per sample", ylab="Number of samples", main="Filtered reads",
     xlim=c(0,4000000), lab=c(10,9,7))
```
