## Supplemental Materials 1 for "Pronounced Genetic Separation Among Varieties of the *Primula cusickiana* Species Complex, a Great Basin Endemic": SNP_Matrix.R

```
# ----Building SNP matrix----
# Script for analyzing the sample by SNP matrix output from ipyrad. 
# The purpose of this script is to determine how many SNPs are shared between replicates, 
# and to also demonstrate how to combine the SNPs found between replicate samples. 
# This script refers to the UniqueReps ipyrad run.

library(vcfR)
setwd("/home/akoontz11/kaiser/Code/ipyrad/UniqueReps/")

# Load in .vcf file using vcfR library
vcf.Object <- read.vcfR("UniqueReps.vcf")
# VCF objects include meta-information, fixed fields, and genotype data. We want the latter.
str(vcf.Object@gt)
# We extract the genotype information, and remove the first column 
# (FORMAT describes the format of all the gt information, which in this example is entirely GT:DP:CATG)
# We also transpose the matrix so that samples are rows, and genotype data are columns
vcf.Object <- t(vcf.Object@gt[,-1])
# Remove DP:CATG data, to only include genotype data (or allele ref.)
vcf.Object <- gsub(pattern=":[0-9]{1,4}:[0-9]{1,4},[0-9]{1,4},[0-9]{1,4},[0-9]{1,4}", replacement="", vcf.Object)

unique(c(vcf.Object))
ncol(vcf.Object)

# Find number of samples (rows) containing an SNP (that is, containing at least one instance that isn't ./.). Sort it.
hits <- apply(vcf.Object,2,function(x) {length(grep(pattern="\\./\\.", x, invert=TRUE))})
hits <- sort(hits, decreasing=TRUE)
hits
x.axis <- 1:ncol(vcf.Object)

# Plot the vector against the number of SNPs (columns)
plot(hits ~ x.axis, xlab="Number of SNPs", ylab="Number of samples per SNP")

# ----Plotting number of SNPs per sample----
SNPs <- apply(vcf.Object,1,function(x) {length(grep(pattern="\\./\\.", x, invert=TRUE))})
# Number of SNPs per sample, including replicates
barplot(height=SNPs, las=3)

reps <- grep(pattern="\\_R", x=names(SNPs))
comb <- SNPs[reps-1] + SNPs[reps]
collapsed_SNPs <- SNPs[-reps]
collapsed_SNPs <- c(collapsed_SNPs[-(reps-1)], comb)
collapsed_SNPs <- collapsed_SNPs[order(names(collapsed_SNPs))]

# Number of SNPs per sample, combining replicates
barplot(height=collapsed_SNPs, las=2, ylab="Number of SNPs", yaxp=c(0,4000,16), main="SNPs per sample, replicates combined")

# ----Determining number of SNPs shared between replicates----
rownames(vcf.Object)
# Ck54 and Ck54_R: 1666 SNPs
a <- grep(pattern="\\./\\.", vcf.Object[24,], invert=TRUE)
length(grep(pattern="\\./\\.", vcf.Object[24,], invert=TRUE))
r <- grep(pattern="\\./\\.", vcf.Object[25,], invert=TRUE)
length(grep(pattern="\\./\\.", vcf.Object[25,], invert=TRUE))
a %in% r
length(which(a %in% r))

# Ck85 and Ck85_R: 1714 SNPs
a <- grep(pattern="\\./\\.", vcf.Object[31,], invert=TRUE)
length(grep(pattern="\\./\\.", vcf.Object[31,], invert=TRUE))
r <- grep(pattern="\\./\\.", vcf.Object[32,], invert=TRUE)
length(grep(pattern="\\./\\.", vcf.Object[32,], invert=TRUE))
a %in% r
length(which(a %in% r))

# Mg40 and Mg40_R: 1653 SNPs
a <- grep(pattern="\\./\\.", vcf.Object[58,], invert=TRUE)
length(grep(pattern="\\./\\.", vcf.Object[58,], invert=TRUE))
r <- grep(pattern="\\./\\.", vcf.Object[59,], invert=TRUE)
length(grep(pattern="\\./\\.", vcf.Object[59,], invert=TRUE))
a %in% r
length(which(a %in% r))

# Mg64 and Mg64_R: 1643 SNPs
a <- grep(pattern="\\./\\.", vcf.Object[66,], invert=TRUE)
length(grep(pattern="\\./\\.", vcf.Object[66,], invert=TRUE))
r <- grep(pattern="\\./\\.", vcf.Object[67,], invert=TRUE)
length(grep(pattern="\\./\\.", vcf.Object[67,], invert=TRUE))
a %in% r
length(which(a %in% r))

# Dm112 and Dm112_R: 1878 SNPs
a <- grep(pattern="\\./\\.", vcf.Object[46,], invert=TRUE)
length(grep(pattern="\\./\\.", vcf.Object[46,], invert=TRUE))
r <- grep(pattern="\\./\\.", vcf.Object[47,], invert=TRUE)
length(grep(pattern="\\./\\.", vcf.Object[47,], invert=TRUE))
a %in% r
length(which(a %in% r))

# Nv134 and Nv134_R: 845 SNPs
a <- grep(pattern="\\./\\.", vcf.Object[82,], invert=TRUE)
length(grep(pattern="\\./\\.", vcf.Object[82,], invert=TRUE))
r <- grep(pattern="\\./\\.", vcf.Object[83,], invert=TRUE)
length(grep(pattern="\\./\\.", vcf.Object[83,], invert=TRUE))
a %in% r
length(which(a %in% r))

# Pa147 and Pa147_R: 1791 SNPs
a <- grep(pattern="\\./\\.", vcf.Object[90,], invert=TRUE)
length(grep(pattern="\\./\\.", vcf.Object[90,], invert=TRUE))
r <- grep(pattern="\\./\\.", vcf.Object[91,], invert=TRUE)
length(grep(pattern="\\./\\.", vcf.Object[91,], invert=TRUE))
a %in% r
length(which(a %in% r))

# Pa152 and Pa158: 21 SNPs
a <- grep(pattern="\\./\\.", vcf.Object[93,], invert=TRUE)
length(grep(pattern="\\./\\.", vcf.Object[93,], invert=TRUE))
r <- grep(pattern="\\./\\.", vcf.Object[96,], invert=TRUE)
length(grep(pattern="\\./\\.", vcf.Object[96,], invert=TRUE))
a %in% r
length(which(a %in% r))

# Pa152 and Pa154: 1956 SNPs
a <- grep(pattern="\\./\\.", vcf.Object[94,], invert=TRUE)
length(grep(pattern="\\./\\.", vcf.Object[94,], invert=TRUE))
r <- grep(pattern="\\./\\.", vcf.Object[96,], invert=TRUE)
length(grep(pattern="\\./\\.", vcf.Object[96  ,], invert=TRUE))
a %in% r
length(which(a %in% r))
```
