## Supplemental Materials 1 for "Pronounced Genetic Separation Among Varieties of the *Primula cusickiana* Species Complex, a Great Basin Endemic": MaguireiOnlyPlot.R

```
# maguirei-only STRUCTURE plot

setwd("/home/austin/Documents/WolfLab/Primula/Analysis/Code/R/structure/")

# %%% READING IN DEMO FILE %%%----
# Reading in K3 file, an updated version of the CLUMPAK K3.output file. 
# Comes from MaguireiOnly ipyrad/STRUCTURE run, .ustr output file from ipyrad
k3 <- read.table("K3_maguirei.csv", header=F)

# Make a vector of names
names <- c("Lower Logan Canyon","Right Hand Fork","Seed Source")
# Make a vector for name positions on graph
labelPos <- c(5, 13, 19)

# Plotting function for K of any value
plot_k <- function(klist,labelPositions,...){
  # List of colors, which are combinations of RGB components, in hexadecimal  
  colors <- c("#2171B5","#D95F02","#7570B3","#E7298A","#66A61E","#8C510A","#666666")
  # Graphing parameters
  par(mar=c(5,2,4,2)+0.1, mgp = c(3,1,1))
  for(i in 1:length(klist)){
    if(i==1){
      # Initial barplot
      barplot(klist[,i], xlim=c(0,20), horiz=F, beside=F, col=colors[i], axisnames=T, space=0.2, yaxt= "n", main="K=3")
      off.value <- klist[,i]
    }else{
      # Subsequent barplots with offset
      barplot(klist[,i], offset=off.value, add=T, beside=F, xlim=c(0,100), horiz=F, col=colors[i], yaxt= "n")
      off.value <- off.value + klist[,i]
    }
  }
  # y axis
  axis(2, at = c(0, 0.25, 0.5, 0.75, 1), labels=c("0", "0.25", "0.50", "0.75", "1.00"), cex.axis = 1, las = 2, pos = -0.2, xpd=T)
  # Add group labels
  text(x=labelPositions, y=-0.03, srt=40, adj=1, xpd=TRUE, labels=names, cex=0.75)
}
dev.off()
plot_k(k3[,2:4], labelPos)

# Plotting lines beneath groups
lineWidth <- 1.3; lineHeight <- rep(-0.015,2)

lines(x = c(0.2,9.6), y = lineHeight, lwd = lineWidth, col = "black", xpd = NA)
lines(x = c(9.8,15.6), y = lineHeight, lwd = lineWidth, col = "black", xpd = NA)
lines(x = c(15.7,21.6), y = lineHeight, lwd = lineWidth, col = "black", xpd = NA)
```
