## Supplemental Materials 1 for "Pronounced Genetic Separation Among Varieties of the *Primula cusickiana* Species Complex, a Great Basin Endemic": multipleKs.R

```
setwd("~/kaiser/Primula/code/structure/JuneSubset/CLUMPAK/")

# load in ks from clummpack files
k2 <- read.csv("K2.csv", sep = '', header = F)
k2 <- k2[,-1]
k3 <- read.csv("K3.csv", sep = '', header = F)
k3 <- k3[,-1]
k4 <- read.csv("K4.csv", sep = '', header = F)
k4 <- k4[,-1]
k5 <- read.csv("K5.csv", sep = '', header = F)
k5 <- k5[,-1]
k6 <- read.csv("K6.csv", sep = '', header = F)
k6 <- k6[,-1]
k7 <- read.csv("K7.csv", sep = '', header = F)
k7 <- k7[,-1]
k8 <- read.csv("K8.csv", sep = '', header = F)
k8 <- k8[,-1]
k9 <- read.csv("K9.csv", sep = '', header = F)
k9 <- k9[,-1]
k10 <- read.csv("K10.csv", sep = '', header = F)
k10 <- k10[,-1]
k11 <- read.csv("K11.csv", sep = '', header = F)
k11 <- k11[,-1]
k12 <- read.csv("K12.csv", sep = '', header = F)
k12 <- k12[,-1]
k13 <- read.csv("K13.csv", sep = '', header = F)
k13 <- k13[,-1]
k14 <- read.csv("K14.csv", sep = '', header = F)
k14 <- k14[,-1]
k15 <- read.csv("K15.csv", sep = '', header = F)
k15 <- k15[,-1]
k16 <- read.csv("K16.csv", sep = '', header = F)
k16 <- k16[,-1]

# Compose large data.frame made up of all columns
x <- as.data.frame(matrix(ncol = 32, nrow = 82))
x[,1:2] <- k2
x[,3:5] <- k3
x[,6:9] <- k4
x[,10:14] <- k5
x[,15:20] <- k6
x[,21:27] <- k7
x[,28:35] <- k8
x[,36:44] <- k9
x[,45:54] <- k10
x[,55:65] <- k11
x[,66:77] <- k12
x[,78:90] <- k13
x[,91:104] <- k14
x[,105:119] <- k15
x[,120:135] <- k16


# lists for plotting
klist_6 <- list(x[,1:2], x[,3:5], x[,6:9], x[,10:14], x[15:20])
klist_12 <- list(x[,21:27], x[,28:35], x[,36:44], x[,45:54], x[55:65])
klist_16 <- list(x[,66:77], x[,78:90], x[,91:104], x[,105:119], x[,120:135])

# Names given to groups
groups <- c("ck_Idaho","ck_Jarbidge","ck_Owyhee","maguirei","domensis","nv_GRBA","nv_Troy", "parryi")

# Matrix for line positions
x.start <- c(0, 25.3, 30.5, 34.8, 53.7, 69.6, 74.7, 78.9)
x.end <- c(25, 30.2, 34.5, 53.4, 69.3, 74.4, 78.6, 86.2)
line.layout <- cbind(x.start, x.end)

dev.off()
structure_plot(ninds = 82, klist_6, groups, line.layout, Ks=2:6)
structure_plot(ninds = 82, klist_12, groups, line.layout, Ks=7:11)
structure_plot(ninds = 82, klist_16, groups, line.layout, Ks=12:16)

#######################################################
# functions needed
######################################################

# plotting and labeling function
structure_plot <- function(ninds = 82, klist, names, L, Ks){
  # define colors (16, for maximum number of Ks)
  cols <- c("#2171B5","#D95F02","#7570B3","#E7298A","#66A61E","#8C510A","#666666","#B3E2CD","#FDCDAC","#CBD5E8","#F4CAE4","#E6F5C9","#FFF2AE","#F1E2CC","#CCCCCC")
  # locations of each column
  b <- as.data.frame(matrix(ncol = 1, nrow = ninds))
  b[,1] <- barplot(t(klist[[1]][1]), beside= F, col= cols, cex.name= 1, cex.axis= 1.2, border = 1, space = 0.05, xaxt = 'n', yaxt = 'n', cex.lab = 1, cex.main=1.6)
  # plot
  plot_q_per_chain(klist, L, Ks)
  # Add labels beneath lowermost plot
  text(cex = 1.5, x = c(13, 26.5, 31, 44, 62, 71, 76, 82), y = -0.275, labels = names, xpd=NA, srt=37, font=3)
}

# plot chains with species labels 
plot_q_per_chain <- function(kqlist, L, Ks, ...){
  # Define colors
  cols <- c("#2171B5","#D95F02","#7570B3","#E7298A","#66A61E","#8C510A","#666666","#B3E2CD","#FDCDAC","#CBD5E8","#F4CAE4","#E6F5C9","#FFF2AE","#F1E2CC","#CCCCCC")
  par(mfrow = c(length(kqlist),1), mar = c(1.5,1,1.3,1) + 0.1, oma = c(5,0,1,0), mgp = c(2,1,0))
  for(i in 1:length(kqlist)){
    # main plot call, and title call
    barplot(t(kqlist[[i]]), beside= F, col= cols, border = 1, space = 0.05, xaxt = 'n', yaxt = 'n', cex.lab = 1.2)
    title(main = paste("K =", Ks[i], sep = ' '), line=0.2, cex.main=1.6)
    # y axis
    axis(2, at = c(0, 0.25, 0.5, 0.75, 1), cex.axis = 1, las = 2, pos = -0.02)
    # lines
    for(j in 1:nrow(L)){
      lines(x = c(L[j,1], L[j,2]), y = c(-0.06,-0.06), lwd = 1.8, col = "black", xpd = NA)
    }
  }
}
```
