## Supplemental Materials 1 for "Pronounced Genetic Separation Among Varieties of the *Primula cusickiana* Species Complex, a Great Basin Endemic": structurePlot.R

```
setwd("/home/austin/Documents/WolfLab/Primula/Analysis/Code/R/structure/")

# %%% READING IN DEMO FILE %%%----
# Reading in K7_subset file, an updated version of the CLUMPAK K7.output file. 
# Comes from JuneSubset ipyrad/STRUCTURE run
# This contains samples in their presented order, and has capillaris and one Troy Peak sample removed from file
k7 <- read.table("K7_plot.csv", header=T)

# Make a vector of names
names <- c("cusickiana_SRP","cusickiana_Jarbidge","cusickiana_Owyhee","maguirei","domensis","nevadensis_GRBA","nevadensis_Troy", "parryi")
# Make a vector for name positions on graph
labelPos <- c(16, 32, 37, 50, 71, 82, 88, 94)

# Plotting function for K of any value
plot_k <- function(klist,labelPositions,...){
  # List of colors, which are combinations of RGB components, in hexadecimal  
  colors <- c("#2171B5","#D95F02","#7570B3","#E7298A","#66A61E","#8C510A","#666666")
  # Graphing parameters
  par(mar=c(5,2,4,1)+0.1, mgp = c(3,1,1))
  for(i in 1:length(klist)){
    if(i==1){
      # Initial barplot
      barplot(klist[,i], xlim=c(0,100), horiz=F, beside=F, col=colors[i], axisnames=T, space=0.2, yaxt= "n", main="K=7")
      off.value <- klist[,i]
    }else{
      # Subsequent barplots with offset
      barplot(klist[,i], offset=off.value, add=T, beside=F, xlim=c(0,100), horiz=F, col=colors[i], yaxt= "n")
      off.value <- off.value + klist[,i]
    }
  }
  # y axis
  axis(2, at = c(0, 0.25, 0.5, 0.75, 1), labels=c("0", "0.25", "0.50", "0.75", "1.00"), cex.axis = 1, las = 2, pos = -0.2, xpd=T)
  # Add group labels
  # text(x=labelPositions, y=-0.026, srt=45, adj=1, xpd=TRUE, labels=names, cex=0.8)
  text(x=labelPositions, y=-0.031, srt=35, adj=1, xpd=TRUE, labels=names, cex=1)
}
dev.off()
plot_k(k7[,4:10], labelPos)
# Plotting lines beneath groups
lineWidth <- 1.9; lineHeight <- rep(-0.015,2)

lines(x = c(0.3,28.3), y = lineHeight, lwd = lineWidth, col = "black", xpd = NA)
lines(x = c(29.4,34.4), y = lineHeight, lwd = lineWidth, col = "black", xpd = NA)
lines(x = c(35.1,39.1), y = lineHeight, lwd = lineWidth, col = "black", xpd = NA)
lines(x = c(40.0,60.7), y = lineHeight, lwd = lineWidth, col = "black", xpd = NA)
lines(x = c(61.5,78.8), y = lineHeight, lwd = lineWidth, col = "black", xpd = NA)
lines(x = c(79.5,84.7), y = lineHeight, lwd = lineWidth, col = "black", xpd = NA)
lines(x = c(85.5,89.5), y = lineHeight, lwd = lineWidth, col = "black", xpd = NA)
lines(x = c(90.5,98.0), y = lineHeight, lwd = lineWidth, col = "black", xpd = NA)

# %%% PLOTTING DIFFERENT CLUSTERS FOR K=6 %%%----
# Code for plotting function
plot_k6 <- function(klist,main,...){
  cols <- c('#A8FFFD','#B862D3', '#A39D9D','#FFFF00', '#69C261', '#FF59AC', '#26CDCD',  '#C1C6FF') # Combination of RGB components, in hexadecimal  
  #par(mfrow = c(length(klist),1), mar=c(0.1,0.1,0.1,0.1)+0.1, oma = c(15,0,0,0), mgp = c(1,1,0))
  for(i in 1:length(klist)){
    if(i==1){
      barplot(klist[,i], xlim=c(0,100), horiz=F, beside=F, col=cols[i], axisnames=T, space=0.2, yaxt= "n", main=main)
      off.value <- klist[,i]
    }else{
      barplot(klist[,i], offset=off.value, add=T, beside=F, xlim=c(0,100), horiz=F, col=cols[i], yaxt= "n")
      off.value <- off.value + klist[,i]
    }
  }
  # y axis
  axis(2, at = c(0, 0.25, 0.5, 0.75, 1), cex.axis = 1, las = 2, pos = -0.2)
}

# Major cluster
k6 <- read.table("ClumppIndFile.output", header=F)
# Remove useless columns
k6 <-k6[,-(1:5)]
plot_k6(k6, main="K=6 Major Cluster")

# Minor cluster 1
k6 <- read.table("ClumppIndFile.MinorCluster1", header=F)
# Remove useless columns
k6 <-k6[,-(1:5)]
plot_k6(k6, main="K=6 Minor Cluster 1")

# Minor cluster 2
k6 <- read.table("ClumppIndFile.MinorCluster2", header=F)
# Remove useless columns
k6 <-k6[,-(1:5)]
plot_k6(k6, main="K=6 Minor Cluster 2")

# %%% DEPRECATED CODE %%%----
# # Read in names file, containing sample IDs, groups, and coordinates (for later use)
# names <- c("Cusickiana_SRP","Cusickiana_Jarbidge","Cusickiana_Owyhee","Maguirei","Domensis","Nevadensis_GRBA","Nevadensis_Troy","Capillaris","Parryi")
# # Vector for name positions
# labelPos <- c(16, 32, 37, 50, 71, 82, 87, 91.5, 97)
# 
# plot_k6 <- function(klist,labelPositions,...){
#   #browser()
#   cols <- c('#A8FFFD','#B862D3', '#A39D9D','#FFFF00', '#69C261', '#FF59AC', '#26CDCD',  '#C1C6FF') # Combination of RGB components, in hexadecimal  
#   #par(mfrow = c(length(klist),1), mar=c(0.1,0.1,0.1,0.1)+0.1, oma = c(15,0,0,0), mgp = c(1,1,0))
#   for(i in 1:length(klist)){
#     if(i==1){
#       barplot(klist[,i], xlim=c(0,100), horiz=F, beside=F, col=cols[i], axisnames=T, space=0.2, yaxt= "n")
#       off.value <- klist[,i]
#     }else{
#       barplot(klist[,i], offset=off.value, add=T, beside=F, xlim=c(0,100), horiz=F, col=cols[i], yaxt= "n")
#       off.value <- off.value + klist[,i]
#     }
#   }
#   # y axis
#   axis(2, at = c(0, 0.25, 0.5, 0.75, 1), cex.axis = 1, las = 2, pos = -0.2)
#   # Add group labels
#   #text(x=labelPositions, y=-0.025, srt=45, adj=1, xpd=TRUE, labels=names, cex=0.75)
# }
# plot.new()
# plot_k6(k6[,3:8], labelPos)
# 
# # Plotting lines beneath groups
# lineWidth <- 1.3; lineHeight <- rep(-0.015,2)
# 
# lines(x = c(0.2,28.5), y = lineHeight, lwd = lineWidth, col = "black", xpd = NA)
# lines(x = c(29,34.7), y = lineHeight, lwd = lineWidth, col = "black", xpd = NA)
# lines(x = c(35.2,39.5), y = lineHeight, lwd = lineWidth, col = "black", xpd = NA)
# lines(x = c(40,61), y = lineHeight, lwd = lineWidth, col = "black", xpd = NA)
# lines(x = c(61.5,79.1), y = lineHeight, lwd = lineWidth, col = "black", xpd = NA)
# lines(x = c(79.6,83.8), y = lineHeight, lwd = lineWidth, col = "black", xpd = NA)
# lines(x = c(84.3,89.8), y = lineHeight, lwd = lineWidth, col = "black", xpd = NA)
# lines(x = c(90.3,92.3), y = lineHeight, lwd = lineWidth, col = "black", xpd = NA)
# lines(x = c(92.8,100.7), y = lineHeight, lwd = lineWidth, col = "black", xpd = NA)

# %%% K6 Stuff %%%

# # Plotting function for K=6
# plot_k6 <- function(klist,labelPositions,...){
#   # List of colors, which are combinations of RGB components, in hexadecimal  
#   cols <- c('#A8FFFD','#B862D3', '#A39D9D','#FFFF00', '#69C261', '#FF59AC', '#26CDCD',  '#C1C6FF') 
#   # Graphing parameters
#   par(mar=c(5,2,4,1)+0.1, mgp = c(3,1,1))
#   for(i in 1:length(klist)){
#     if(i==1){
#       # Initial barplot
#       barplot(klist[,i], xlim=c(0,100), horiz=F, beside=F, col=cols[i], axisnames=T, space=0.2, yaxt= "n", main="K=6")
#       off.value <- klist[,i]
#     }else{
#       # Subsequent barplots with offset
#       barplot(klist[,i], offset=off.value, add=T, beside=F, xlim=c(0,100), horiz=F, col=cols[i], yaxt= "n")
#       off.value <- off.value + klist[,i]
#     }
#   }
#   # y axis
#   axis(2, at = c(0, 0.25, 0.5, 0.75, 1), labels=c("0", "0.25", "0.50", "0.75", "1.00"), cex.axis = 1, las = 2, pos = -0.2, xpd=T)
#   # Add group labels
#   text(x=labelPositions, y=-0.025, srt=45, adj=1, xpd=TRUE, labels=names, cex=0.8)
# }
# plot.new()
# plot_k6(k6[,3:8], labelPos)
# # Plotting lines beneath groups
# lineWidth <- 1.3; lineHeight <- rep(-0.015,2)

# Names with capillaris
# # Read in names file, containing sample IDs, groups, and coordinates (for later use)
# names <- c("cusickiana_Idaho","cusickiana_Jarbidge","cusickiana_Owyhee","maguirei","domensis","nevadensis_GRBA","nevadensis_Troy","capillaris","parryi")
# # Vector for name positions
# labelPos <- c(16, 32, 37, 50, 71, 82, 87, 91.5, 97)

# k6 <- read.table("K6.output", header=F)
# # Remove useless columns
# k6 <-k6[,-(2:5)]
# # Reorder columns based on demo file
# k6 <- k6[k2$Ind,]
# # Give sample names
# k6 <- cbind(k2$Group, k6)
# k6[,3:8]
```
