## Supplemental Materials 1 for "Pronounced Genetic Separation Among Varieties of the *Primula cusickiana* Species Complex, a Great Basin Endemic": README.md

# Overview

This repository contains the scripts and parameter files used for the genomic analysis of the <em>Primula cusickiana</em> species complex. 
It is continuously being updated as the manuscript is in preparation. 

## ipyrad 
The `ipyrad` directory contains 3 folders, each corresponding to a unique run of the [ipyrad](https://ipyrad.readthedocs.io/en/latest/) software pipeline. Directories contain the relevant `params` files used for each run, as well as relevant outfiles (although not all output files are included, as this is unecessary and would require too much space).

The `PreliminaryRuns` directory contains the ipyrad barcode and parameters files used for initial exploratory run (titled "UniqueReps"), along with output files. `ComplexWide` holds the same files for the final run including all complex member samples, and `MaguireOnly` contains the relevant ipyrad files for the run examining only maguirei samples.

In addition, `ComplexWide` and `MaguireiOnly` include folders which contain the files used for F<sub>ST</sub> calculations (using [VCFtools](https://vcftools.github.io/index.html)).

## R
The `R` directory contains the R script used in data analysis. This includes the plotting code for STRUCTURE plots (in `structure`) and DAPC (in `DAPC`). Code used to generate Jaccard distribution of samples is included in the `JaccardDistances.R` script. 

Additionally, included are R scripts used in intial data exploration.

## Mapping

The `Mapping` directory includes the Python script and .csv file used to generate the map of samples with the cluster membership at STRUCTURE output K=7. Credit goes to Dr. Carol Rowe for compiling this Python script!

## STRUCTURE
The `structure` directory contains folders holding the `mainparams` and `extraparams` files for two STRUCTURE runs: the complex wide analysis, and the analysis looking at strictly maguirei samples. Additionally, output files (.str, .stru, and .vcf) are included, as well as the BASH script used to run STRUCTURE.

# Notes
For any questions regarding these scripts, please contact Austin Koontz (either through GitHub or via email). Other scripts/parameter files used in the analysis of this dataset can be made available upon request.
Raw data has been uploaded to the NCBI short-read archive (SRA), accession number PRJNA705310.
