## Supplemental Materials 1 for "Pronounced Genetic Separation Among Varieties of the *Primula cusickiana* Species Complex, a Great Basin Endemic": extraparams.txt

EXTRA PARAMS FOR THE PROGRAM structure.  THESE PARAMETERS CONTROL HOW THE
PROGRAM RUNS.  ATTRIBUTES OF THE DATAFILE AS WELL AS K AND RUNLENGTH ARE 
SPECIFIED IN mainparams.

"(int)" means that this takes an integer value.
"(d)"   means that this is a double (ie, a Real number such as 3.14).
"(B)"   means that this variable is Boolean 
        (ie insert 1 for True, and 0 for False).

PROGRAM OPTIONS

#define NOADMIX     0 // (B) Use no admixture model (0=admixture model, 1=no-admix)
#define LINKAGE     0 // (B) Use the linkage model model 
#define USEPOPINFO  0 // (B) Use prior population information to pre-assign individuals
                             to clusters
#define LOCPRIOR    0 //(B)  Use location information to improve weak data

#define FREQSCORR   1 // (B) allele frequencies are correlated among pops
#define ONEFST      0 // (B) assume same value of Fst for all subpopulations.

#define INFERALPHA  1 // (B) Infer ALPHA (the admixture parameter)
#define POPALPHAS   0 // (B) Individual alpha for each population
#define ALPHA     1.0 // (d) Dirichlet parameter for degree of admixture 
                             (this is the initial value if INFERALPHA==1).

#define INFERLAMBDA 0 // (B) Infer LAMBDA (the allele frequencies parameter)
#define POPSPECIFICLAMBDA 0 //(B) infer a separate lambda for each pop 
					(only if INFERLAMBDA=1).
#define LAMBDA    1.0 // (d) Dirichlet parameter for allele frequencies 


PRIORS

#define FPRIORMEAN 0.01 // (d) Prior mean and SD of Fst for pops.
#define FPRIORSD   0.05  // (d) The prior is a Gamma distribution with these parameters

#define UNIFPRIORALPHA 1 // (B) use a uniform prior for alpha;
                                otherwise gamma prior
#define ALPHAMAX     10.0 // (d) max value of alpha if uniform prior
#define ALPHAPRIORA   1.0 // (only if UNIFPRIORALPHA==0): alpha has a gamma 
                            prior with mean A*B, and 
#define ALPHAPRIORB   2.0 // variance A*B^2.  


#define LOG10RMIN     -4.0   //(d) Log10 of minimum allowed value of r under linkage model
#define LOG10RMAX      1.0   //(d) Log10 of maximum allowed value of r
#define LOG10RPROPSD   0.1   //(d) standard deviation of log r in update
#define LOG10RSTART   -2.0   //(d) initial value of log10 r

                         
USING PRIOR POPULATION INFO (USEPOPINFO)

#define GENSBACK    2  //(int) For use when inferring whether an indiv-
                         idual is an immigrant, or has an immigrant an-
                         cestor in the past GENSBACK generations.  eg, if 
                         GENSBACK==2, it tests for immigrant ancestry 
                         back to grandparents. 
#define MIGRPRIOR 0.01 //(d) prior prob that an individual is a migrant 
                             (used only when USEPOPINFO==1).  This should 
                             be small, eg 0.01 or 0.1.
#define PFROMPOPFLAGONLY 0 // (B) only use individuals with POPFLAG=1 to update	P.
                                  This is to enable use of a reference set of 
                                  individuals for clustering additional "test" 
                                  individuals.

LOCPRIOR MODEL FOR USING LOCATION INFORMATION

#define LOCISPOP      0    //(B) use POPDATA for location information 
#define LOCPRIORINIT  1.0  //(d) initial value for r, the location prior
#define MAXLOCPRIOR  20.0  //(d) max allowed value for r


OUTPUT OPTIONS

#define PRINTNET     1 // (B) Print the "net nucleotide distance" to screen during the run
#define PRINTLAMBDA  1 // (B) Print current value(s) of lambda to screen
#define PRINTQSUM    1 // (B) Print summary of current population membership to screen

#define SITEBYSITE   0  // (B) whether or not to print site by site results. 
		     	       (Linkage model only) This is a large file!
#define PRINTQHAT    0  // (B) Q-hat printed to a separate file.  Turn this 
                           on before using STRAT.
#define UPDATEFREQ   100  // (int) frequency of printing update on the screen.
                                 Set automatically if this is 0.
#define PRINTLIKES   0  // (B) print current likelihood to screen every rep
#define INTERMEDSAVE 0  // (int) number of saves to file during run

#define ECHODATA     1  // (B) Print some of data file to screen to check
                              that the data entry is correct.
(NEXT 3 ARE FOR COLLECTING DISTRIBUTION OF Q:)
#define ANCESTDIST   0  // (B) collect data about the distribution of an-
                              cestry coefficients (Q) for each individual
#define NUMBOXES   1000 // (int) the distribution of Q values is stored as 
                              a histogram with this number of boxes. 
#define ANCESTPINT 0.90 // (d) the size of the displayed probability  
                              interval on Q (values between 0.0--1.0)


MISCELLANEOUS

#define COMPUTEPROB 1     // (B) Estimate the probability of the Data under 
                             the model.  This is used when choosing the 
                             best number of subpopulations.
#define ADMBURNIN  500    // (int) [only relevant for linkage model]: 
                             Initial period of burnin with admixture model (see Readme)
#define ALPHAPROPSD 0.025 // (d) SD of proposal for updating alpha
#define STARTATPOPINFO 0  // Use given populations as the initial condition 
                             for population origins.  (Need POPDATA==1).  It 
                             is assumed that the PopData in the input file 
                             are between 1 and k where k<=MAXPOPS.
#define RANDOMIZE      1  // (B) use new random seed for each run 
#define SEED        2245  // (int) seed value for random number generator 
	                     (must set RANDOMIZE=0) 
#define METROFREQ    10   // (int) Frequency of using Metropolis step to update
                             Q under admixture model (ie use the metr. move every
                             i steps).  If this is set to 0, it is never used.
                             (Proposal for each q^(i) sampled from prior.  The 
                             goal is to improve mixing for small alpha.)
#define REPORTHITRATE 0 //   (B) report hit rate if using METROFREQ
