## Supplemental Materials 1 for "Pronounced Genetic Separation Among Varieties of the *Primula cusickiana* Species Complex, a Great Basin Endemic": mainparams.txt

KEY PARAMETERS FOR THE PROGRAM structure.  YOU WILL NEED TO SET THESE
IN ORDER TO RUN THE PROGRAM.  VARIOUS OPTIONS CAN BE ADJUSTED IN THE
FILE extraparams.


"(int)" means that this takes an integer value.
"(B)"   means that this variable is Boolean 
        (ie insert 1 for True, and 0 for False)
"(str)" means that this is a string (but not enclosed in quotes!) 


Basic Program Parameters

#define MAXPOPS    2      // (int) number of populations assumed
#define BURNIN    50000   // (int) length of burnin period
#define NUMREPS   100000   // (int) number of MCMC reps after burnin

Input/Output files

#define INFILE   infile   // (str) name of input data file
#define OUTFILE  outfile  //(str) name of output data file

Data file format

#define NUMINDS    82    // (int) number of diploid individuals in data file
#define NUMLOCI    1227    // (int) number of loci in data file
#define PLOIDY       2    // (int) ploidy of data
#define MISSING     -9    // (int) value given to missing genotype data
#define ONEROWPERIND 0    // (B) store data for individuals in a single line


#define LABEL     1     // (B) Input file contains individual labels
#define POPDATA   0     // (B) Input file contains a population identifier
#define POPFLAG   0     // (B) Input file contains a flag which says 
                              whether to use popinfo when USEPOPINFO==1
#define LOCDATA   0     // (B) Input file contains a location identifier

#define PHENOTYPE 0     // (B) Input file contains phenotype information
#define EXTRACOLS 0     // (int) Number of additional columns of data 
                             before the genotype data start.

#define MARKERNAMES      0  // (B) data file contains row of marker names
#define RECESSIVEALLELES 0  // (B) data file contains dominant markers (eg AFLPs)
                            // and a row to indicate which alleles are recessive
#define MAPDISTANCES     0  // (B) data file contains row of map distances 
                            // between loci


Advanced data file options

#define PHASED           0 // (B) Data are in correct phase (relevant for linkage model only)
#define PHASEINFO        0 // (B) the data for each individual contains a line
                                  indicating phase (linkage model)
#define MARKOVPHASE      0 // (B) the phase info follows a Markov model.
#define NOTAMBIGUOUS  -999 // (int) for use in some analyses of polyploid data


Command line options:

-m mainparams
-e extraparams
-s stratparams
-K MAXPOPS 
-L NUMLOCI
-N NUMINDS
-i input file
-o output file
-D SEED
