## Supplemental Materials 1 for "Pronounced Genetic Separation Among Varieties of the *Primula cusickiana* Species Complex, a Great Basin Endemic": MaguireiOnly.sh

```
#!/bin/bash
for k in {2..8} ;
do
    for r in {1..50} ;
    do
        $HOME/apps/structure/console/structure  -i MaguireiOnly.str -m mainparams.txt -e extraparams.txt -K $k -o output/MaguireiOnly_output_$k-$r > output/outfile_$k_$r  &
        sleep 3s
    done
done
echo "All runs started"
```
